# How we draw and recognize things that don’t exist

**DOI:** 10.64898/2026.04.22.720107

**Authors:** Emily J. A-Izzeddin, Filipp Schmidt, Christian Houborg, Henning Tiedemann, Roland W. Fleming

## Abstract

How do we make sense of something we’ve never seen before? Classifying objects into superordinate classes like ‘animal’ is a key step in interpreting novel experiences, but is challenging because radically different items (e.g., octopus, rabbit) must somehow be grouped together. In general, no single feature is shared by all members. We reasoned that to classify or imagine novel items from outside the distribution of previous experiences, observers parse objects into meaningful component features that they can mentally recombine (‘compositionality’). To test this, we asked participants to draw familiar and novel members of nine superordinate object classes. We then asked other participants to classify the drawings, and mark and label their defining ‘parts’. We find that human classification performance is well predicted by a Bayesian classifier that optimally combines the part labels, suggesting humans can create and classify out-of-distribution experiences through a compositional generative representation of object features.

## 1. Introduction

The ability of the human mind to classify objects into distinct categories is fundamental for making sense of the world, guiding our behaviour, and facilitating effective communication and learning^1,2^. By treating different objects as equivalent, we organise a complex and varied world into meaningful groups that share common characteristics, allowing us to make prompt decisions based on prior experiences and knowledge. For young children first learning about the world, a key challenge is working out what kind of entity a completely novel stimulus might be (e.g., a previously unencountered animal, plant, or tool). Yet, while there is a considerable body of research on the recognition and classification of familiar objects^3,4^, how we perceive and understand things we have never seen before (i.e., zero-shot learning) remains poorly understood. Identifying novel items requires generalising outside the distribution of previous experience, presumably through an internal model of the typical features associated with each class that can be recombined in novel configurations. Here, we used creative drawings—by asking people to draw pictures of new, non-existent kinds of objects—to probe their internal models. Testing how other people group these pictures into familiar categories tells us how we make sense of things we have never seen before.

### 1.1 The difficulty of grouping of objects into classes

Objects can be classified at different levels of abstraction. The *subordinate* level is most precise, with objects grouped into highly specific classes with specific features (e.g., North Pacific Giant Octopus). The *basic* level is less specific, with objects sharing common features (e.g., octopus). The superordinate level is the most general, grouping objects into broad, overarching classes (e.g., animal)^5^. This continuum of abstraction organises information hierarchically.

The actual level of classification used in any given scenario varies depending on expertise, behavioural goals, or the variety of encountered samples^6^. The basic level is often considered to be psychologically privileged as it provides enough information for meaningful classification while also being specific enough to be useful in everyday contexts^7^.

However, when we encounter an object for the first time, the most important distinction is often at the superordinate level (e.g. animal, plant, or tool) as this determines the range of possible responses. Indeed, the first time we encounter something, it is, by definition, out of distribution (‘OOD’) at the basic level, so superordinate classification is the best hope we have of working out what it might be. Yet, superordinate classification is also arguably the most enigmatic, given that objects with often vastly different visual appearances must somehow be grouped together. For example, octopuses, rabbits, and hummingbirds are all animals (**Fig. 1**), yet share few features in common. Somehow, we must abstract out the combination of features that gives an animal its typical ‘look and feel’. It remains unclear how humans solve this challenge.

**Figure 1.**
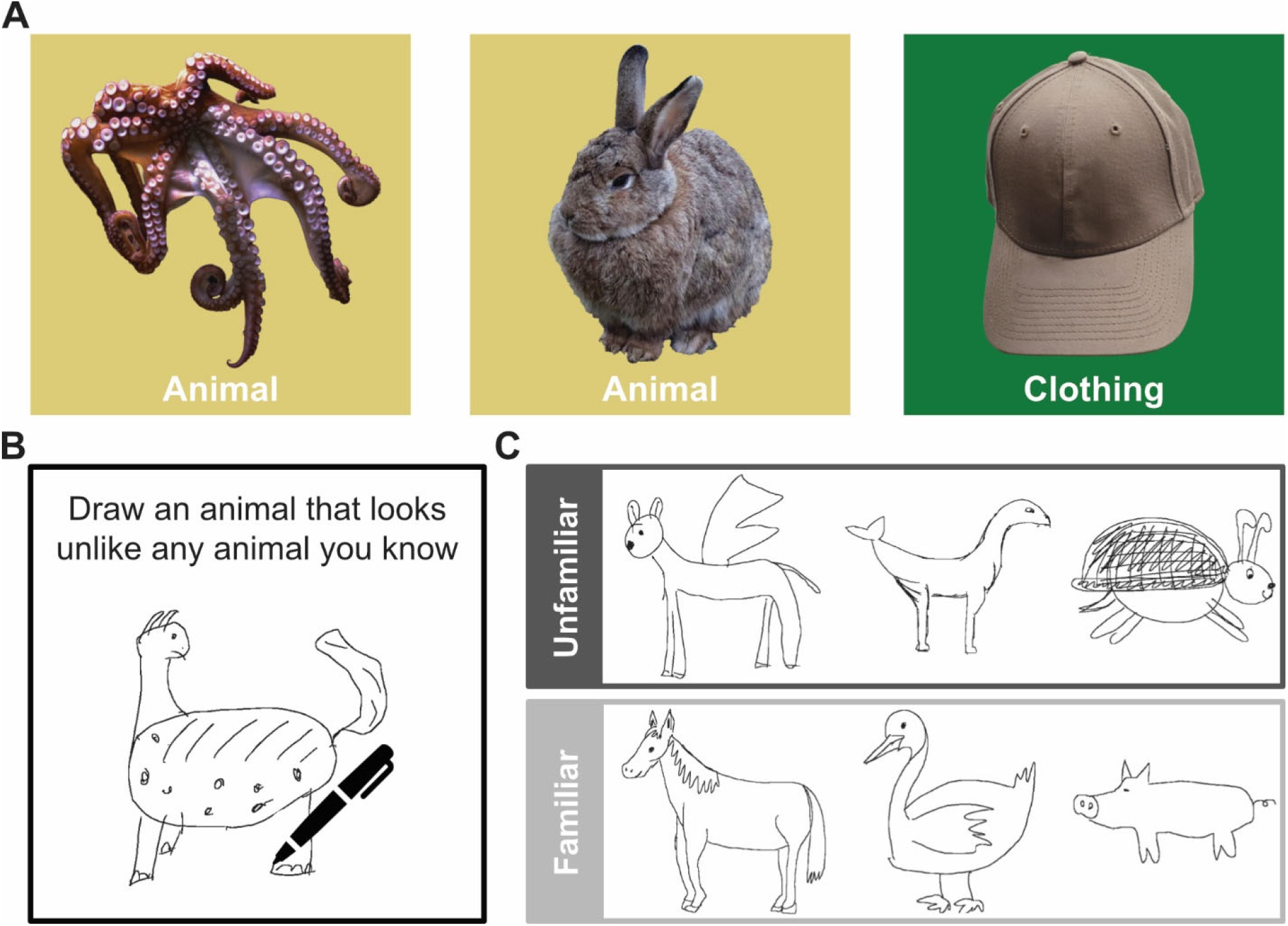
The challenge of superordinate classification. **A)** Appearance can vary substantially within a class (e.g. animals), to such an extent that overall shape and appearance might be more different within a class (octopus–rabbit) than between classes (rabbit–cap). **B)** We had participants generate drawings of novel unfamiliar object examples for specific superordinate classes (here ‘animal’). For comparison, we also did the same for familiar objects. **C)** Examples of unfamiliar (top row) and familiar (bottom row) drawings generated as examples from the ‘animal’ superordinate class.

### 1.2 Discriminative vs generative approaches to classification and the power of compositionality

A major open question is whether our ability to make such distinctions reflects a straightforward ‘discriminative’ model (e.g., feedforward processes based on embedding of stimuli relative to class boundaries in feature space), or alternatively, a more sophisticated ‘generative’ model that would allow explicit top-down re-sampling to imagine what novel exemplars might look like. This question gets at the heart of a fundamental debate on the role of discriminative and generative models in perception and cognition^8^. A key hypothesis is that the features we use to parse and represent objects are not arbitrary—which would allow discrimination but make the creation of novel examples challenging—but instead reflect the semantic ontology of the real world (e.g., eyes, head, legs). This would allow ‘compositionality’^9–11^, that is, the ability to represent combinations of features that have not been encountered before, both to classify and synthesise novel objects. We reasoned that rather than have people simply observe, discriminate, and classify *familiar* objects, a more fruitful way to probe for the existence and nature of generative models is to have people explicitly create (i.e., draw) new examples of things that *do not exist* and therefore require OOD generalisation. We suggest the easiest way for participants to succeed at this task (i.e., to create things that others would agree belong to the desired superordinate class) would be to recombine semantically meaningful features associated with the classes.

### 1.3 Challenges to understanding superordinate classification of novel objects

Previous studies of superordinate classification have largely focused on animals, suggesting classification depends on certain diagnostic features (e.g., eyes, mouth, limbs)^3^, which to some extent predict the division of animate and inanimate objects in behavioural responses and neural activity^12^. Yet it is unclear how well this applies to other superordinate classes. Moreover, focussing on familiar objects conflates visual appearance and semantic knowledge about those samples. For example, when seeing the image of a cow, we can not only identify individual appearance features (e.g., eyes, horns, udder, boxy shape, etc.) but also identify it as a cow, making it difficult to distinguish the relative importance of bottom-up and top-down information flow.

There have been attempts to tease apart visual appearance and semantic knowledge. For example, in a previous study^13^, we presented observers with ambiguous stimuli (like the famous duck-rabbit shape^14^) along with a word to specify which interpretation they should adopt, and found that the interpretation of the local features of the shape changed radically based on the class (i.e., a top-down effect). We also investigated the perception of stimuli that are misclassified at the superordinate level (e.g., sea-anemone, which is an animal that is routinely perceived as a plant by nonexperts). The responses hinted at constellations of mid-level, statistical shape features (e.g., symmetry, curvature) that drove both correct and incorrect classifications^15^. Another study found neural representations in early visual cortex to better reflect visual appearance features (e.g., head, mouth, eyes) than animacy when comparing images of animals and lookalike inanimate objects^16^. However, the features driving superordinate classification were not specifically investigated. Other studies have modified images by various scrambling or morphing processes to prevent objects from being explicitly recognised while conserving certain mid-level appearance features^17–20^. Yet this approach may eliminate the very high-level visual appearance features like “head” and “mouth” that are most important in our day-to-day classifications of real objects.

Previous findings therefore seem to hint at visual appearance features being important for superordinate classification, at least for animals. However, it remains unclear to what extent visual appearance features facilitate superordinate classification more broadly, spanning a wider range of superordinate classes, and especially with unfamiliar (novel) examples. Here, we aimed to overcome some limitations of previous studies—conflation of appearance and semantic knowledge, experimenter bias in picking relevant features, and the absence of truly novel samples with high-level appearance features—by asking participants to draw novel, out-of-distribution samples of superordinate classes.

### 1.4 Using generative drawings to acquire out-of-distribution samples

How do we use drawings to identify the mechanisms underlying superordinate object classification for newly encountered objects? We reason that if humans have a detailed mental representation of object classes, they should be able to generate novel members of these classes, and drawing provides a practical approach to testing this. Consequently, drawing has received increasing attention in cognition and perception research as a method to directly probe mental concepts^21,22^. For example, studies have used drawings to demonstrate memory fidelity and spatial biases for visual scenes^23^, individual differences in internal representations for typical scene arrangements^24^, as well as humans’ capacity for generating previously unencountered examples of simple^25^ and complex shapes^26,27^. Drawings therefore pose an opportunity to probe humans’ internal representations for the key features that are definitive of a particular superordinate class of objects, and how these may be recombined to form novel members of a given class.

Here, we had participants draw novel members of nine superordinate classes: animals, buildings, clothing, furniture, household appliances, musical instruments, plants, tools, and vehicles. We then asked other participants to assess the drawings’ class and typicality, as well as identify and label the drawings’ defining ‘parts’. Importantly, by asking participants to generate non-existent objects, we prompted them to draw instances of class members that fall into the relevant area of their representational space—i.e., expresses significant features that communicate the intended superordinate class (e.g., of animals)—without referring to specific members of that class (e.g., cow). In doing so, we reduce the conflation of visual appearance features and semantic information related to known objects. We then tested whether defining parts labelled by observers were sufficient to predict the perceived class. While no individual part was especially diagnostic, our findings indicate a near-optimal (Bayesian) use of part combinations, shared by the creators and observers of the drawings. Together our findings suggest observers have rich generative internal models of superordinate object classes, based on a compositional coding of semantic features, which enables both the creation and zero-shot recognition of novel, OOD stimuli.

## 2. Results

### 2.1 Observers successfully generate unfamiliar members of familiar classes

We asked participants to draw either novel (Experiment 1A; **Fig. 2**) or familiar (Experiment 1B; **Fig. 2**) members of cued superordinate classes.

**Figure 2.**
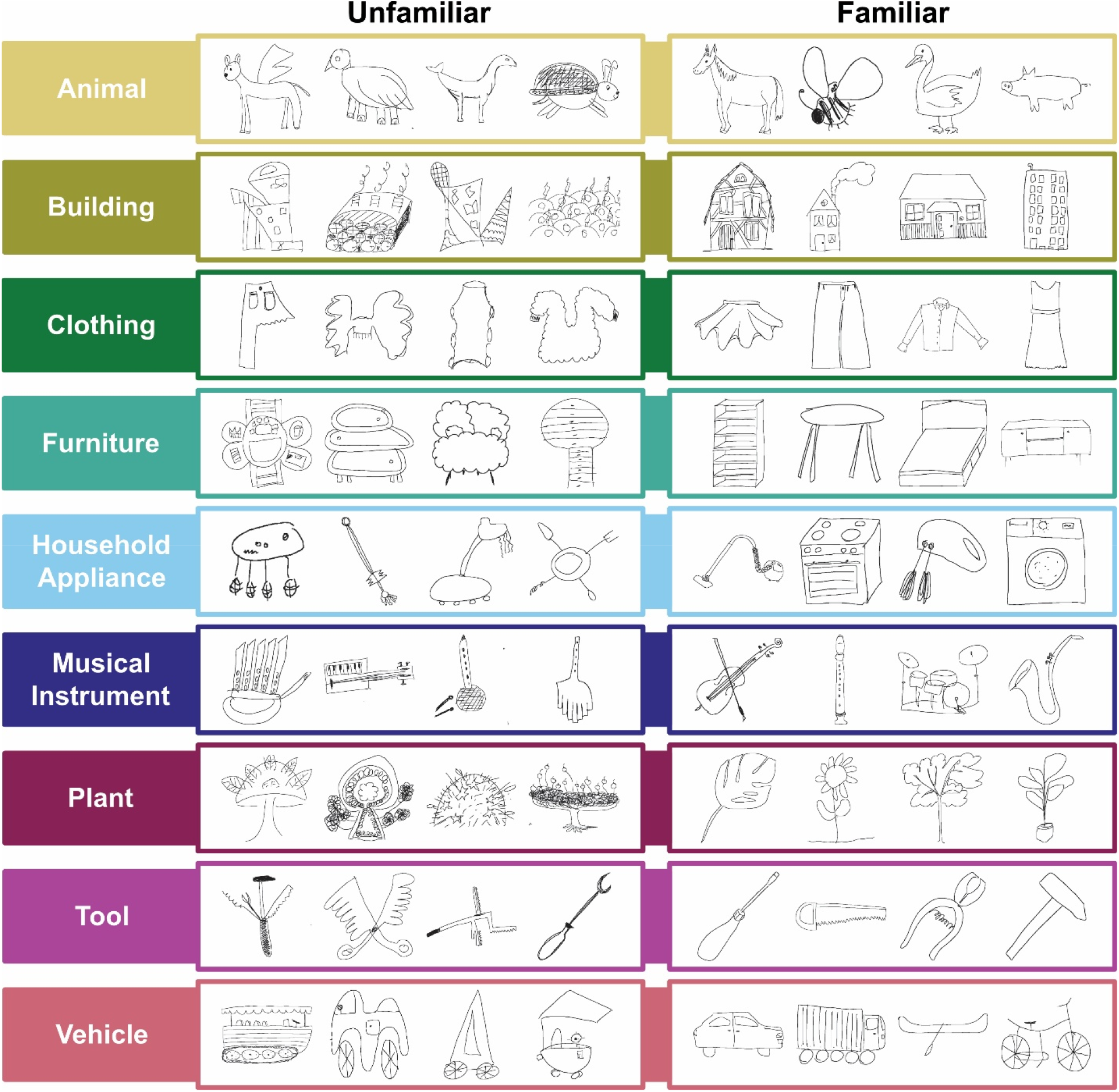
Participant drawings. Example drawings from Experiment 1A (unfamiliar, left) and Experiment 1B (familiar, right), separated by instructed superordinate class (rows). Note: Object drawings are not included as they were not analysed further for the familiar drawings.

To assess whether the drawings were both novel and classifiable, we had an independent group of participants judge the drawings’ typicality and superordinate class (Experiment 2; **Fig. 3A**). They could select out of the nine prompted classes or ‘other’. We tested the proportion of times each drawing was identified as the same superordinate class that was cued for the original drawers. This revealed extreme evidence for classification performance to be well above chance (10%) for both unfamiliar (overall *m* = 62%, *sd* = 0.38, one sample t-test *BF*_10_ = 2.818×10^183^, *t* = 39.06, *p* < .001) and familiar (overall *m* = 89%, *sd* = 0.23, *BF*_10_ = 2.000×10^47^, *t* = 32.21, *p* < .001) drawings (**Fig. 3B**).

**Figure 3.**
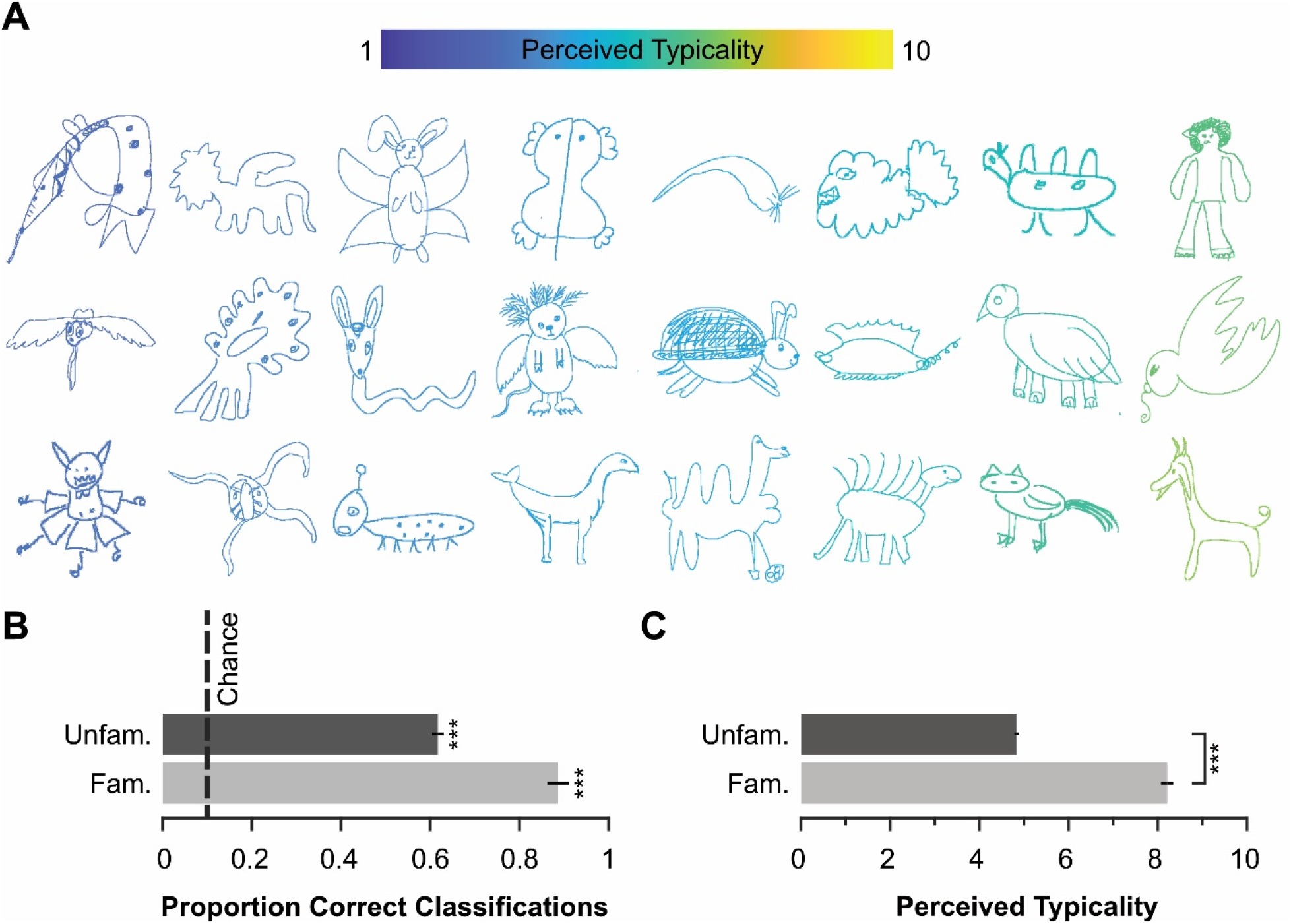
Classification and typicality ratings for generated drawings. **A)** Example unfamiliar animal drawings from Experiment 1A, arranged and colour-coded according to their average perceived typicality. **B)** The average proportion of correct classification of images (x-axis) for unfamiliar and familiar drawings (y-axis) relative to chance (dotted line). *** *BF*_10_ > 10 in favour of a difference existing between chance and classification performance. **C)** Same as B, but now showing average typicality ratings. *** *BF*_10_ > 10 in favour of a difference in perceived typicality existing between familiar and unfamiliar drawings. Error bars represent +/- 1 SEM.

This suggests the drawers were effective at generating classifiable members of known superordinate classes.

Crucially, however, we were interested in whether participants in Experiment 1A were effective at generating novel, *unfamiliar* members of the classes. If they were successful, the perceived typicality of their unfamiliar drawings should be lower than that of familiar drawings (**Fig. 3A**; all 800 unfamiliar and 90 familiar drawings are plotted in the **Supplemental Materials, Figs. S1-11**, and are available for download at https://osf.io/bn63v/overview?view_only=284fa8175f4544bb81aacfd8f07bdfb2). We find very strong evidence for typicality (rated on a scale from 1-10) being rated much lower for unfamiliar drawings (overall *m* = 4.84, *sd* = 1.53) than for familiar drawings (overall *m* = 8.21, *sd* = 1.33, independent samples t-test *BF*_10_ = 1.660×10^70^, *t* = 20.062, *p* < .001; **Fig. 3C**). Taken together, drawers appear to have been very effective at generating drawings of novel, but classifiable members of known superordinate classes.

The validity of the drawings is further supported by their embeddings in the latent space of the DreamSim model^28^, which uses both low-level image features and semantic/categorical inferences to predict perceptual similarities between images. We used t-SNE^29^ to visualise the spread and localisation of these embeddings in 2D (**Fig. 4**). We found little difference in the spread of unfamiliar and familiar drawings when considering both the average pairwise distances of drawings within each class (paired sample t-test *BF*_10_ = 0.363, *t* = -0.62, *p* = .550), as well as the mean distance of each image to its class’ centroid (*BF*_10_ = 1.070, *t* = -1.85, *p* = .098). Indeed, when we compare the pairwise distances between the centroids of each class for familiar and unfamiliar drawings, we find a strong correlation (*r* = .88, *BF*_10_ = 6.716×10^12^, *p* < .001; **see Section 4.4.4 for analysis details**). Therefore, it seems that unfamiliar and familiar drawings yield similar spatial relationships in their clustering.

**Figure 4.**
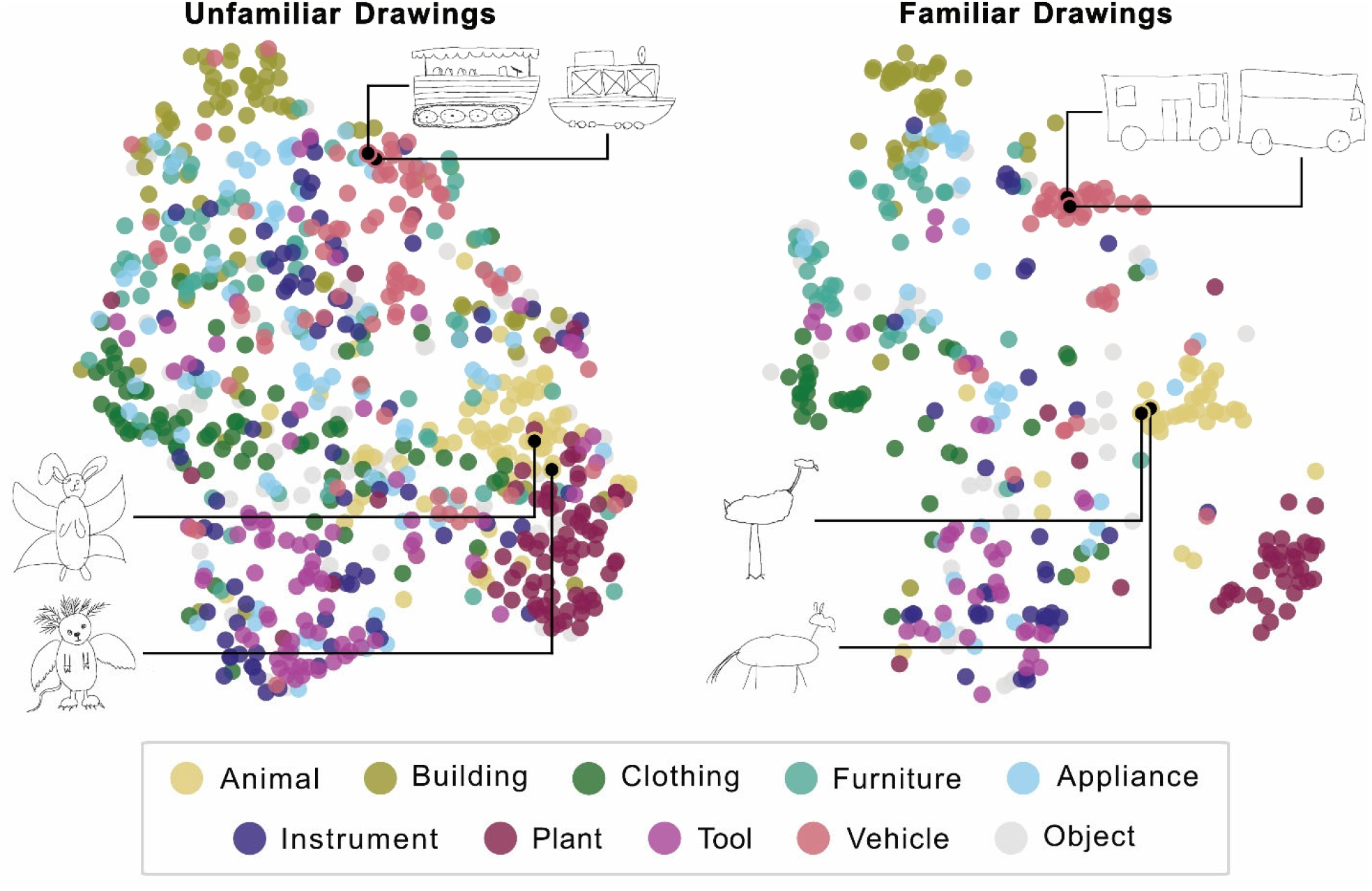
DreamSim embeddings for unfamiliar and familiar drawings. Here, we plot a two-dimensional representation of the DreamSim embeddings for both unfamiliar (left) and familiar (right) drawings, acquired through t-distributed Stochastic Neighbor Embedding (t-SNE). Points are colour-coded according to the cued superordinate class, as indicated by the legend. In addition, two examples of vehicles and animals are highlighted for both plots, demonstrating the effectiveness of DreamSim at grouping similar drawings together, regardless of whether the objects are unfamiliar or familiar. NOTE: all 400 familiar drawings are plotted here to allow improved interpretability of the DreamSim output, rather than the sub-sample of 90 drawings.

Additionally, we inspected the relationship between typicality ratings and classification accuracy for unfamiliar drawings. Overall, we found very strong evidence for a moderate positive correlation between typicality ratings and classification performance (*r* = .37, *BF*_10_ = 2.123×10^24^, *p* < .001). At the individual class level, correlations were much weaker for all familiar drawings, as well as unfamiliar animals and buildings, due to ratings being heavily skewed towards high typicality/high classification accuracy (see **Supplemental Materials, Figs. S12-13**, for more detail). Nonetheless, our results suggest better classification performance for drawings that are perceived to be more typical. Such a relationship follows given that an unfamiliar drawing must inherently have been encountered less often (or, in this case, never), necessitating their status as atypical to the viewer, as compared with familiar drawings.

### 2.2 The majority of drawing strokes are grouped into distinct parts

To investigate the features relevant for superordinate classification, we had other participants mark and label the relevant ‘parts’ of drawings that drove their classification (Experiment 3; example in **Fig. 5A**). Both familiar and unfamiliar drawings were far above zero in terms of the number of parts labelled (one sample T-test minimum *BF*_10_ = 3.362×10^32^, *t* = 20.74, *p* < .001) and the portion of pixels labelled (minimum *BF*_10_ = 1.288×10^84^, *t* = 87.45, *p* < .001). We found that unfamiliar drawings, on average, had 2.47 parts labelled (*sd* = 0.96), and familiar drawings had 3.02 parts labelled (*sd* = 1.38; **Fig. 5C**). There was very strong evidence for this difference (independent samples T-test *BF*_10_ = 9119.473, *t* = 4.86, *p* < .001), which likely follows from participants’ knowledge about specific object parts and their associated labels in familiar objects. At the same time, the proportion of pixels labelled was 0.67 for unfamiliar drawings (*sd* = 0.10) and 0.76 (*sd* = 0.08) for familiar drawings (**Fig. 5D**). There was very strong evidence for this difference (independent samples T-test *BF*_10_ = 2.364×10^13^, *t* = 8.38, *p* < .001), again, likely due to participants’ greater understanding of the parts present in familiar drawings and their boundaries. Nonetheless, for both familiar and unfamiliar drawings, on average the majority of pixels are labelled, suggesting the drawings are made up of strokes attributable to recognisable parts that observers report being relevant to their superordinate classifications.

**Figure 5.**
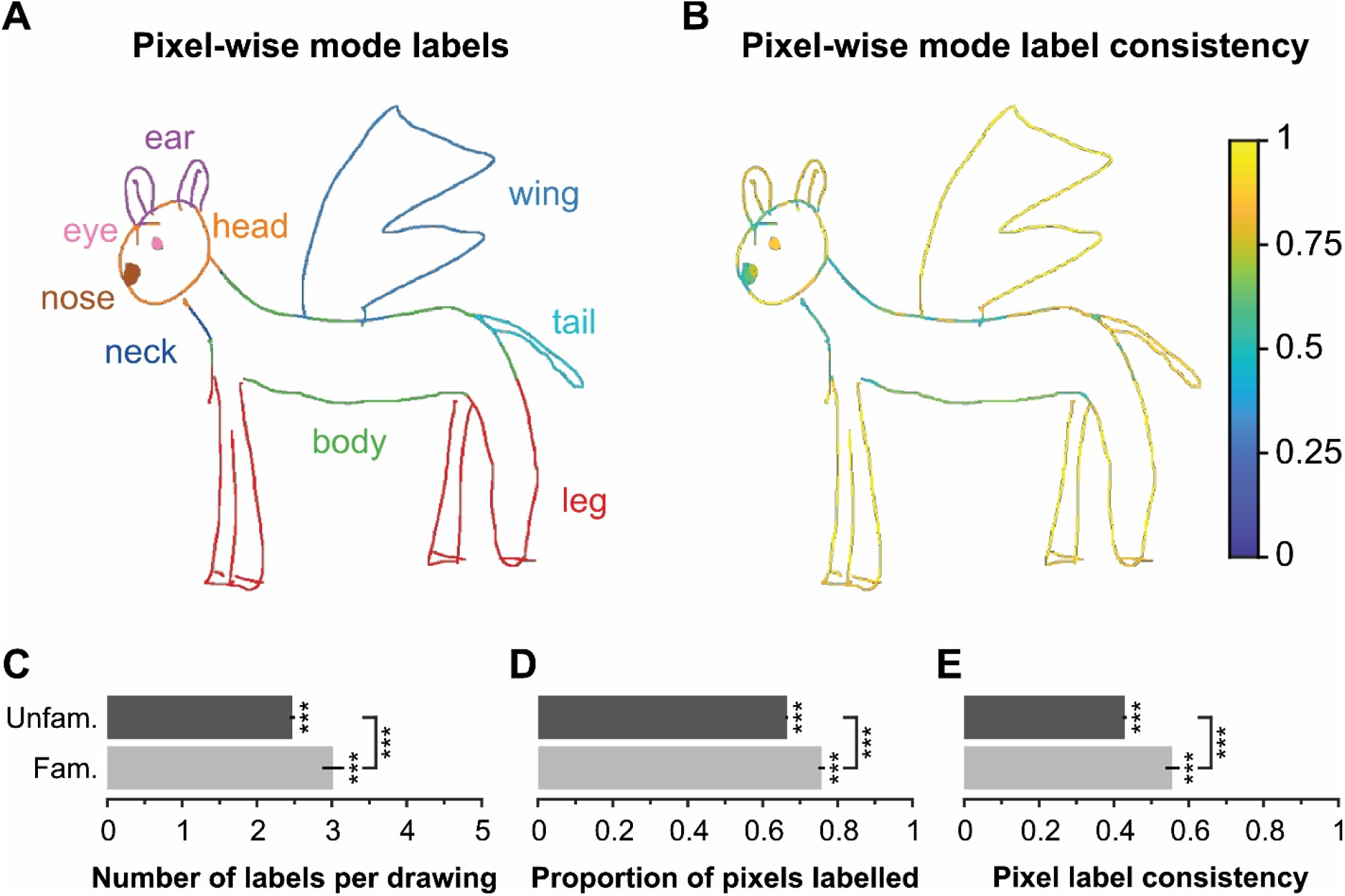
The calculation of pixel-wise consistency. **A)** Example marking data for one drawing, showing the pixel-wise mode labels across participants’ responses, colour-coded by unique mode label. **B)** Pixel-wise consistency of the mode labels for the same drawing, ranging from 0 to 1 (where 1 indicates all observers reported the same label for a given pixel). **C)** The average number of labels per drawing (x-axis) for unfamiliar and familiar drawings (y-axis). **D)** Same as **C**, however now plotting the average proportion of drawing pixels labelled. **E)** Same as **C**, however now plotting the average pixel-wise label consistency. Error bars represent +/- 1 SEM. *** *BF*_10_ > 1,000 in favour of a difference existing between familiar and unfamiliar drawings.

### 2.3 Even unfamiliar drawings are made up of recognisable parts

We next sought to examine how recognisable parts were, particularly for unfamiliar drawings. To do so, we determined the pixel-wise consistency of responses – i.e., for a given pixel, we identified the mode label and the proportion of observers who provided that mode label (**Fig. 5A-B**). This allows us to identify regions of a given drawing that yielded relatively consistent labels and calculate the overall labelling consistency for each drawing.

To identify the overall consistency of labels for a given drawing, we simply take the average consistency score for all pixels within that drawing. We found very strong evidence for greater consistency for familiar drawings (*m* = 0.56, *sd* = 0.18) than unfamiliar drawings (*m* = 0.43, *sd* = 0.17, independent samples t-test *BF*_10_ = 4.297×10^8^, *t* = 6.82, *p* < .001; **Fig. 5E**). This likely again reflects observers’ inherent and shared interpretation of the drawn objects and their parts. Nonetheless, both familiar and unfamiliar drawings were labelled with consistency scores well above zero (one-sample T-test minimum *BF*_10_ = 1.950×10^44^, *t* = 29.55, *p* < .001).

### Drawings in superordinate classes yield distinct, but sparse part labels

We sought to identify the critical elements or ‘parts’ of an object that facilitate superordinate classifications. As participants marked and labelled distinct parts of the drawings (Experiment 3), part labels were attributed to specific pixels of specific drawings, presenting many potential avenues for analyses. Firstly, we were interested in gaining a basic impression of the types of labels (reflecting the parts present in the drawings), and whether these differ between superordinate classes (see **Fig. 6A**). To look at the label distinctiveness between superordinate classes, we performed an intersection over union (IoU) analysis. For a given IoU calculation, we take the unique labels given for two superordinate classes, calculate the number of shared labels between the two classes, and divide this by the total number of unique labels across both classes. The resulting value ranges from 0-1, where 0 indicates there is no overlap, and 1 indicates complete overlap. We did this in a pairwise manner across all possible superordinate class combinations, as defined by the classification made by participants in Experiment 3 who also labelled the parts, and inspected only the upper triangle values in the resulting matrices. This analysis revealed very little overlap in labels between superordinate classes for both familiar (mean pairwise IoU = 0.07, *sd* = 0.04) and unfamiliar drawings (*m* = 0.13, *sd* = 0.05), with values significantly below 1 in both cases (one sample T-test minimum *BF*_10_ = 1.492×10^53^, *t* = 12.27, *p* < .001; see **Fig. 6B**). Also, we find very strong evidence for a positive correlation between the IoU matrices for unfamiliar vs familiar drawings (*r* = .79, *BF*_10_ = 1.185×10^8^, *p* < .001), suggesting consistency in the distinctiveness of labels between superordinate classes regardless of familiarity.

**Figure 6.**
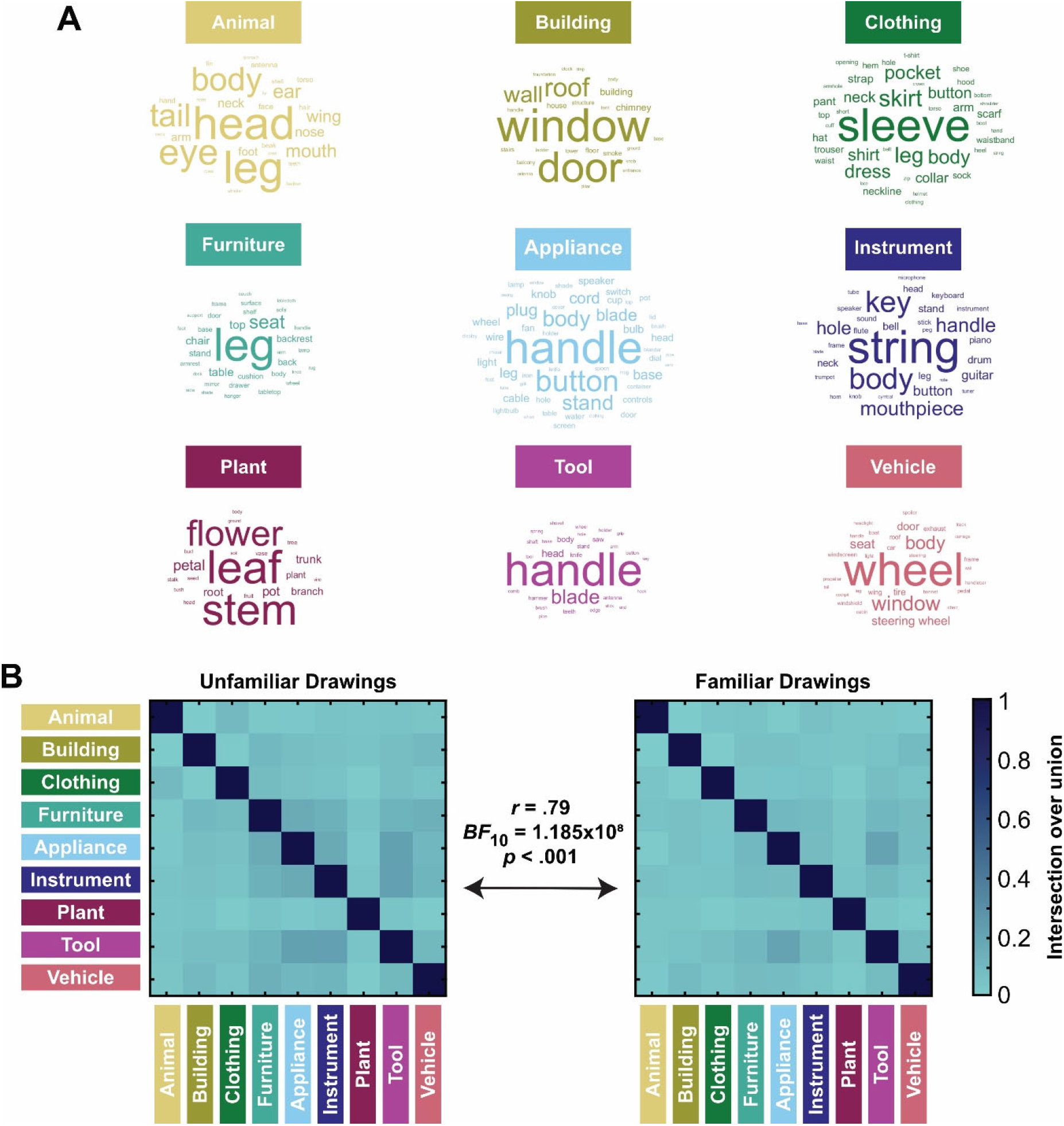
Parts associated with different superordinate classes. **A)** Word clouds yielded from the part labels provided for unfamiliar drawings. Labels vary in size to represent relative frequency. Labels are only included if they are applied to more than 1% of drawings within the class. **B)** The results of an intersection over union analysis completed for superordinate class labels given for unfamiliar (left) and familiar (right) drawings. Here, superordinate class is defined by the classification given by participants in Experiment 3.

We thought that the distinctiveness of labels might be related to the ability of participants to perform accurate classifications. To investigate this, we correlate the IoU score for each superordinate class pairing (per **Fig. 6**) with the number of classification confusions specific to each pairing. As expected, for unfamiliar drawings (excluding the negative diagonal, representing correct classifications), we find very strong evidence for a moderate positive correlation (*r* = .52, *BF*_10_ = 113503.515, *p* < .001), demonstrating more classification confusions when the IoU score is greater (for more detail and figures, see the **Supplemental Materials, Figs. S14-15**). Taken together, our results suggest distinct parts are of use to observers when classifying drawn objects into superordinate classes. This is further supported by more frequent misclassifications between classes sharing parts.

In addition to the distinctiveness *between* superordinate classes, we were also interested in the degree of overlap in part labels *within* superordinate classes. We therefore calculated the IoU on a trial-wise basis, allowing comparison of labels given for different images within the same superordinate class. We then took the average IoU score for each superordinate class combination to replicate the matrices in **Fig. 6B**. In doing so, we can inspect the sparseness of labels within a class, with high IoU scores indicating high consistency of parts identified across class members, and low scores indicating high variability in the identified parts. Overall, we find quite low IoU scores within superordinate classes. For unfamiliar drawings, the maximum IoU score within a single class was only .19 (buildings), and for familiar drawings was .30 (buildings; **see Supplemental Materials, Fig. S16, for further visualisation**), suggesting most individual drawings only present a small subset of the parts associated with their class. Furthermore, we observe very few parts that appear in a majority of trials attributed to a single superordinate class. For example, for unfamiliar drawings, only three parts were present on more than half of trials attributed to the relevant superordinate class. These were: “wheel” within the class of vehicles (63%), “window” within the class of buildings (59%), and “door” within the class of buildings (53%). Indeed, for the vast majority of superordinate classes, across both familiar and unfamiliar drawings, the frequency of the most common label could not account for the classification accuracy observed (exception: “leg” label for unfamiliar furniture drawings was present on 42% of trials).

These results therefore suggest no single characteristic or part is sufficient to define a superordinate class. Instead, members are related to one another by ‘family resemblance’^30^. Indeed, sparse part sets are likely necessary to capture the enormous variability of features possible for radically different objects within a single superordinate class (**Fig. 1A**). Thus, while each superordinate class yields distinct labels (**Fig. 6B**), it appears unlikely for one object to possess the complete set of features necessary to define a single superordinate class. Such sparsity in features within a superordinate class further highlights our impressive aptitude for superordinate classification.

### 2.5 Superordinate classification aligns with Bayesian label combination

Given the sparse, varied set of parts required to define a single superordinate class, we aimed to assess the extent to which we could account for participants’ classifications based on such labels. Specifically, we hypothesised that participants inform their classifications via some form of optimal combination of individual parts and their respective diagnosticity for the possible superordinate classes. We therefore implemented a Bayesian classification approach wherein, for a given image presented to a given participant, we combine the likelihoods of the individual parts identified. From here, we calculate the posterior for each possible superordinate class and assume the class with the highest posterior probability is selected (for more details, see **Section 4.4.3**). We completed this classification on a trial-wise basis, allowing direct comparison of a Bayesian classifier to observers’ judgements.

We trained the classifier on the priors and part likelihoods for *familiar* drawings and then tested how well the classifier predicted the perceived class of the *unfamiliar* drawings. On a trial-by-trial basis, we found a close correspondence between observer and classifier responses of 65%, well above the chance level of 10% (binomial test *BF*_10_ = ∞, *p* < .001). Indeed, we found extreme evidence for a positive correlation between participant confusions (**Fig. 7A**) and classifier confusions (*r* = .90, *BF*_10_ = 2.511×10^33^, *p* < .001; **Fig. 7B;** see **Supplemental Materials, Fig. S18**, for classifier confusion matrix). As not all part labels present for unfamiliar drawings were also present for familiar drawings, some unfamiliar drawings had no likelihood information to derive from their parts. In this case, the classification is driven by the class priors, which are based on the number of drawings classified into the various superordinate classes by participants. Such ratings resulted in a higher count of animal objects than any other class, meaning that in the case where no likelihood information can be derived from the parts, the classifier selected animal. As a result of this bias, some data points in **Figure 7B** fall away from the diagonal. The high correspondence between participants’ classifications and those of a Bayesian classifier suggests human superordinate classifications can be largely explained by an optimal combination of image parts. Further, such an optimal combination appears to be sufficient for mitigating the difficulty posed by the sparsity of parts that define superordinate classes.

**Figure 7.**
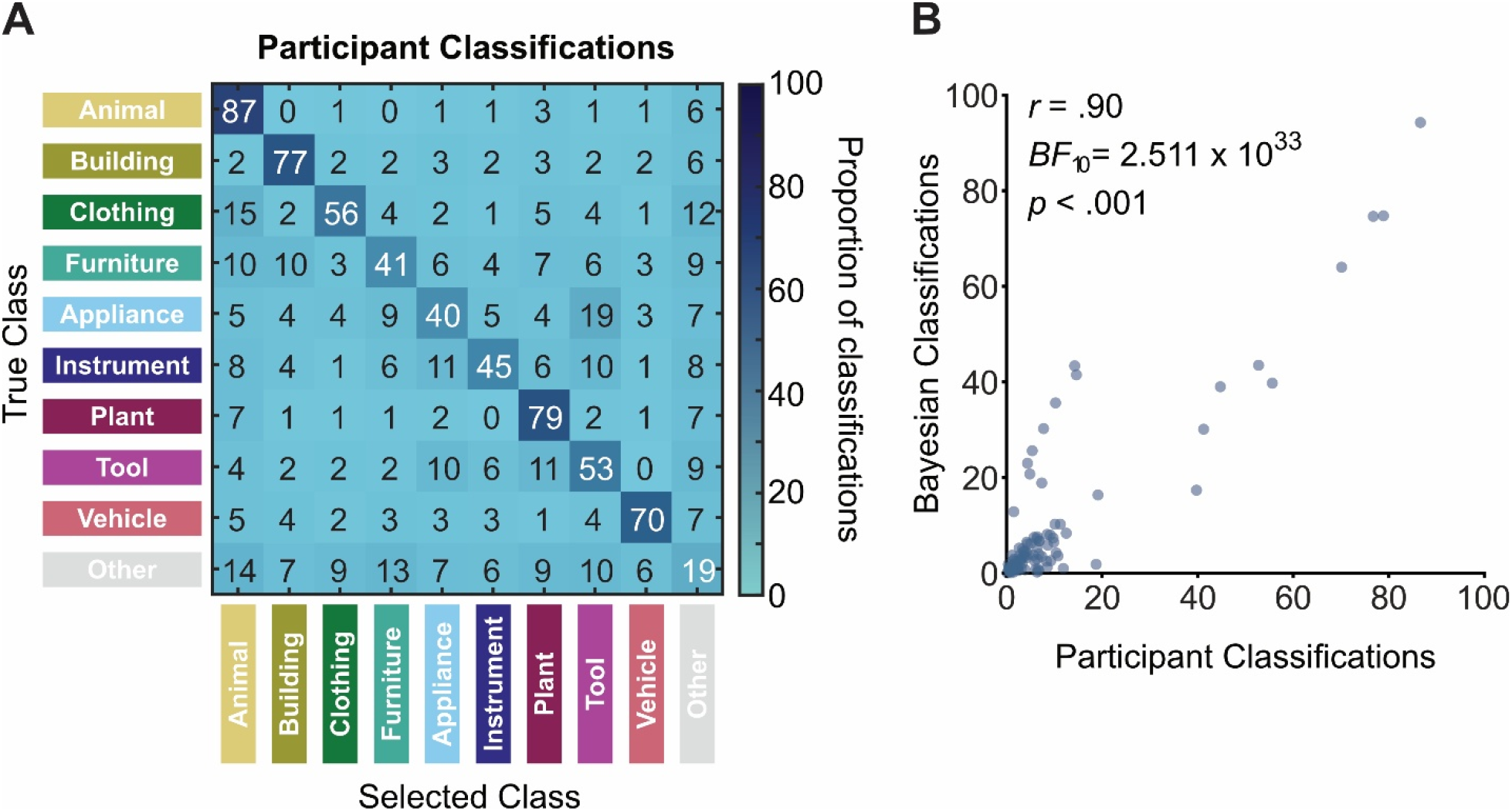
Human superordinate classifications reflect near-optimal part combination. **A)** Matrix representing the percentage of responses where an unfamiliar drawing belonging to a given superordinate class (rows) was classified as another given class (columns), representing the number of classification confusions (with the exception of the negative diagonal, representing correct classifications). Due to rounding, some rows do not perfectly add up to 100. **B)** The relationship between classification confusions made by participants (Panel A, including the negative diagonal), and those made by a Bayesian classifier for unfamiliar drawings which bases its classifications on prior/likelihood information from familiar drawings.

## 3. Discussion

We set out to investigate how humans perceive and understand things we have never seen before. We hypothesised that this requires generalising outside the distribution of previous experience and is presumably accomplished through an internal generative model of the typical features associated with each class, which can be recombined in novel configurations. To test this idea, we asked a group of participants to draw novel exemplars of objects from nine superordinate classes. We then asked other observers to classify the drawings and identify and label the parts in each drawing that drove their classification.

### 3.1 Humans can generate drawings of novel but interpretable class members

We found that participants can invent novel members of superordinate classes, for example, making up genuinely new animals, tools, or clothing. This was demonstrated by consistently lower typicality ratings for novel vs familiar drawings by the other participants. Humans therefore seem to have a rich mental representation of object classes, which they can tap into to generate novel members from a wide range of classes, creating a multitude of novel object appearances. Our results therefore extend previous findings related to humans’ ability to generate novel shape exemplars, demonstrating that we can indeed do so for far more complex and varied object classes^26^. Importantly, these were not random, unrecognisable scribbles. On the contrary, they were highly structured so that independent observers were generally very good at relating the novel drawings to their intended, familiar class. This suggests humans are not only effective in object recognition but also in visually communicating object classes, even at the superordinate level. Generating and identifying such novel items inherently requires us to generalise outside the distribution of our previous experiences, but how do we do this?

### 3.3 Important parts vary significantly both between and within classes

We hypothesised that the most effective means of communicating superordinate classes would be by including particular parts associated with the class (e.g. wheels, to signify a vehicle). Adults only need two to four major parts to recognise instances of common classes^31,32^, and even young children classify objects based on the parts they perceive to be shared among objects. For example, Rakison and Butterworth (1998) found that 1- to 2-year-old children show intuitive classifications of animals and objects based on shared parts, such as legs and wheels^33^. Also, 18- to 30-month-old children can match novel objects based on the shape of parts (despite differences in part relations), but not vice versa – suggesting they organise objects into parts and mainly use those, rather than arrangement, to guide their classification judgements of novel objects^34^.

To test our hypothesis, we had observers mark and label the individual parts (e.g., head, window, petal) that lead to their classification. From these labels, we could interrogate the distinctiveness of nameable parts between and within classes. We found that there was very little overlap in the parts labelled between classes, suggesting largely unique sets that define different superordinate classes. And even though the scope of our study is much wider, where previous evidence is available – as for the classification of familiar animals – it is largely consistent with our findings (eyes, mouth, limbs in animacy perception)^3,12^.

However, notably we also found large variability of labels *within* each class, suggesting there are no single parts capable of comprehensively defining a superordinate class. This reflects the extraordinary degree of diversity between members of a single superordinate class and explains why even though distinctive parts help to classify objects into superordinate classes, this is still a major computational challenge when object identity is unknown. It is therefore far from trivial that creators and observers of the drawings shared an understanding of what type of object was being depicted. Moreover, the high consistency between participants in terms of which regions of the images they labelled as distinctive parts—and their ability to give them semantically meaningful labels—further supports the notion of a compositional representation of superordinate classes.

### 3.4 Bayesian inference can largely account for superordinate classification

The sparse and varied sets of parts present between and within superordinate classes inherently prevents single parts from being sufficiently diagnostic of a given class. We therefore hypothesised that superordinate classification may follow from the combination of parts present for a given drawing (cf. Wittgenstein’s notion of ‘family resemblance’^30^). To test whether observers combine features optimally, we implemented a Bayesian classifier which groups drawings based on the evidence provided by the set of its parts. We found this model yielded remarkably consistent classification results to those of our observers, suggesting a near-optimal use of parts in the generation and classification of novel objects. This is remarkable, given that previous work often emphasised the importance of mid-level features, such as curvature or symmetry for unfamiliar animals^15,17–19^. Our current results suggest classification can also be attributed to much more semantically-based mechanisms, in terms of nameable parts. Such a contribution of semantically-relevant features supports the notion of a generative compositional framework for classification, whereby humans can flexibly represent combinations of features that have not been encountered previously.

Of course, the current task inherently promotes emphasis of the nameable areas in observers’ representational space—indeed, the use of line drawings inherently limits the types of information that can be used to depict an object. For example, colour has been shown to play some role in object classification^12,35–37^, even though this might be limited to colour-diagnostic objects^38^ and be less important for some classes like animals^3^. In future studies it would be interesting to allow drawers to use colour, or indeed other features like varying line weight. We might also expect a contribution of texture or material appearance (e.g., fur). Even though it is possible to render complex textures or material appearances in drawings, our (naïve) participants might have been biased by the experienced difficulty with which certain features can be drawn. Note, however, that we included drawings of typical objects as a control (which are subject to the same limitations)^21^. Further, our results nonetheless suggest superordinate classification of novel objects *can* be performed and accounted for by an optimal combination of known features.

Notably, Bayesian label combination, as implemented here, ignores the number and spatial organisation of features within an image, which are both potentially important characteristics of superordinate classes (e.g., the number of legs and their positioning in animals vs furniture). It therefore remains unclear to what extent classification performance is dependent on such factors. Even though our current model already predicts human behaviour very well, future studies might use stimuli and models that include more object features and spatial organisation of exemplars.

### 3.5 Conclusions

We investigated the mechanisms underlying generation and superordinate classification for previously unencountered, novel objects. To do so, we used participant-generated drawings to probe humans’ representational space for superordinate classes by asking them to generate novel members of known classes. We had other observers classify the drawings and mark and label the parts they felt were most indicative of the drawings’ class. We posited that people would both create and interpret the drawings in a *compositional* manner by relying on novel combinations of semantically meaningful features associated with each class. Observers were indeed successful at generating genuinely novel, but identifiable drawings. From the provided part labels, we found that those used to generate both familiar and novel objects vary greatly between superordinate classes, promoting their distinctiveness. However, we found that superordinate classes cannot be captured by any single part but, instead, are necessarily characterised by a large superset of potential parts, with only a few of which featured in any given instance. We suggest this was necessary to effectively represent the enormous variability of objects within each superordinate class. Critically, despite the significant computational challenge presented by such within-class diversity, we found that the classifications were highly consistent with a Bayesian classifier. In other words, novel image generation and classification can be explained by near-optimal use of feature combinations. Together, our results suggest observers have rich, generative, and compositional internal models of superordinate classes and their constituent semantic features, enabling the creation and zero-shot recognition of novel, OOD stimuli.

## 4. Methods

### 4.1 Funding

This study was funded by the European Research Council (project number 101098225 - ERC-2022-AdG “STUFF”), by the European Union’s Horizon Europe research and innovation programme under the Marie Skłodowska-Curie grant agreement No. 101226908 (‘EXPLORA’), and by the Deutsche Forschungsgemeinschaft (German Research Foundation, DFG, project number 222641018–SFB/TRR 135 Project C1) as well as under Germany’s Excellence Strategy (EXC 3066/1 “The Adaptive Mind”, Project No. 533717223).

### 4.2 Open practices statement

All data and stimuli are available at https://osf.io/bn63v/overview?view_only=284fa8175f4544bb81aacfd8f07bdfb2. This study’s design and its analysis were not pre-registered.

### 4.3 Participants and procedures

Experiments were approved by the Local Ethics Committee of the Department of Psychology and Sports Sciences of the Justus Liebig University Giessen (LEK-2020-0041 and LEK-2021-0032). Participants were recruited via a university mailing list (Experiments 1-2) or Prolific (Experiment 3). Participants in all experiments had self-reported normal or corrected-to-normal vision and were naïve to the purpose of the study. They gave informed consent and were treated in accordance with the ethical guidelines of the American Psychological Association (APA). All participants received financial compensation or course credits for their participation.

#### 4.3.1 Experiment 1A: Drawing novel class members

In the first experiment, participants were asked to draw novel, unfamiliar members of different superordinate classes. Specifically, the instructions that were shown above the drawing area always had the form of “Draw a plant that looks unlike any plant you know” (translated from German: “Zeichne eine Pflanze, die aussieht wie keine Dir bekannte Pflanze”). The classes probed were: animals, buildings, clothing, furniture, household appliances, musical instruments, plants, tools, and vehicles (chosen based on their coverage in two public datasets of object images and drawings, THINGS+^39^ and QuickDraw^40^). Per class, each participant produced 5 drawings consecutively, after which the prompt changed to a different class. The order of classes was randomised. An additional ‘object’ condition was included at the end, where participants were asked to “Draw an object that looks unlike any object you know”.

Participants drew on a 12.9” touchpad in portrait orientation using a touchpad pen. The drawing area was 14 by 14 cm in size. There was no time limit and participants could draw as much as they wanted. Overall, 16 participants took part in the experiment, creating 800 drawings (see left-hand column of drawings in **Fig. 2** for examples).

#### 4.3.2. Experiment 1B: Drawing typical class members

For comparison with the novel drawings, we asked a new group of participants (N = 8) to draw 5 typical, familiar class members for each of the classes, resulting in 400 familiar drawings (see right-hand column of drawings in **Fig. 2** for examples). The drawing interface was identical to the previous experiment.

#### 4.3.3 Experiment 2: Measuring classification and typicality

To measure to what extent the drawings can be considered legitimate members of their classes and their level of novelty, we conducted a rating task with a new group of participants (n = 32). For Experiment 2, all drawings from Experiment 1A and 90 drawings from Experiment 1B (10 drawings from each class, except from the ‘object’ condition) were used. The full image dataset was split into two sets of 445 drawings, balanced across classes and including both novel and typical drawings, with each set being completed by 16 participants. Drawings were shown individually in randomised order. In each trial, participants were asked first to assign the drawing to a class, selecting one of the 9 class labels or an “other” option which allowed them to type in their response. Second, they were asked to rate how typical that drawing was for its class by setting a slider from 1 (typical) to 10 (not typical).

#### 4.3.4 Experiment 3A: Identifying drawing parts for unfamiliar drawings via human observers

To identify the defining features for each unfamiliar drawing, we conducted a marking task with a new group of participants (n = 160 after replacing exclusions, see **Section 4.4.1** for exclusion details). Here, all drawings from Experiment 1A were shown. Each participant viewed 50 unique drawings from the set individually, presented in randomised order, with each drawing being observed by 10 different participants. The sequence of assigned drawings to be viewed was generated by repeatedly shuffling and assigning drawings until all participants were assigned 50 unique drawings and each drawing had been assigned 10 times. In each trial, participants were first asked to classify the drawing into one of 10 classes (the same as in Experiment 1A) (Instruction: “First, select the category you think the sketch belongs to.”). Participants were asked to avoid using the “other” option where possible, and to provide their own label when choosing “other”. After classifying a drawing, participants marked and labelled a defined ‘part’ of the drawing that contributed to their choice of class (“Afterwards, you will be asked to highlight the different parts of the sketch that lead you to provide your chosen category.”). To define parts, they coloured in drawing strokes using the cursor and then provided a written label for that part. Participants then repeated this procedure for the same drawing until they felt all defined parts of a drawing were marked – participants were required to mark at least one part, and a maximum of 15 parts. Finally, they could proceed to the next drawing.

#### 4.3.5 Experiment 3B: Identifying drawing parts for familiar drawings via human observers

Here, we applied the same general approach as in Experiment 3A, but for the familiar drawings (the same 90 drawings from Experiment 1B as rated in Experiment 2). A new set of participants were recruited (N = 20 after replacing exclusions, see **Section 4.4.1** for exclusion details). Each participant viewed 40 or 50 drawings (due to counterbalancing). Otherwise, procedures were identical to Experiment 3A, with each drawing observed by 10 different participants.

### 4.4 Data processing

#### 4.4.1 Participant exclusions and data cleaning

Experiments 1-2 (sub-experiments included) did not involve any specific participant exclusions or data cleaning – the following pertains only to Experiment 3 (i.e., part marking/labelling). A technical error resulted in two participants in Experiment 3A completing trial assignments that had already been completed by existing participants. To avoid uneven numbers of data points for each drawing, we simply removed the repeat participants’ data from analyses. In addition, one participant in Experiment 3A provided a blank label on 26 of the 50 trials. This participant was removed from the dataset and their trial data was replaced by a new participant.

After the completion of data collection, we performed checks to identify any participants who needed to be excluded due to poor task performance. Our first exclusion check measured the proportion of pixels marked by the participants. Outliers were identified as participants who, on average, marked less than two standard deviations below the mean. This resulted in the exclusion of nine participants in Experiment 3A and two participants in Experiment 3B. We also included two checks of the data to identify whether or not participants had followed instructions. Firstly, we identified participants who marked the drawing but input multiple labels per part despite being instructed to provide single part labels. Secondly, we identified participants who frequently provided superordinate class labels as part labels (e.g. ‘animal’). Participants were excluded if more than 30% of their responses fell into either of the above categories: 14 participants in Experiment 3A, and one participant in Experiment 3B. All excluded participants’ data were replaced by new participants, resulting in final sample sizes of 160 for Experiment 3A and 20 for Experiment 3B.

Once we had complete datasets (after replacing exclusions), basic processing of the provided labels was completed to remove unnecessary noise from the data. A script was used to perform basic cleaning of the labels: all letters were converted to lowercase, punctuation (i.e., commas, periods, slashes, double spaces, parentheses, etc.) was removed, spaces at the beginning or end of a label were removed, all labels were converted to plurals, and any labels with multiple words were converted to single words (i.e., by simply adjoining the multiple words) given that the resulting single-word form was already present amongst the complete list of unaltered labels. Following automated processing, two independent raters went through the entire list of unique labels, converted them back to singular, removed adjectives, corrected obvious spelling mistakes (in cases of uncertainty, labels were left unaltered), and converted all spelling to British English. Any labels containing more than one part in the label were changed to refer only to the first listed. Finally, any labels in which participants provided nonwords or explicitly stated they did not know what label to apply were converted to missing values, with that part removed from the marking data. Because label cleaning was completed independently of viewing the associated drawings (and marked parts of said drawings), the intended meaning of particular labels was often ambiguous – as such, we did not attempt to homogenise synonyms amongst the labels. Original label data was replaced with cleaned labels for the purposes of analyses. Finally, when cleaning resulted in identical labels for different parts of the same drawing from a single participant, they were combined into one part; which reduced the total number of labelled parts across drawings and participants from 20,063 to 19,778 for unfamiliar drawings (0.01% reduction), and from 2,733 to 2,715 for familiar drawings (0.01% reduction).

#### 4.4.2 General analysis procedure

All frequentist statistics were performed in MATLAB (version R2022b), while all Bayesian analyses were completed in JASP (version 0.19.3). Input data for inferential statistical analyses comprise the mean responses for each drawing (i.e., averaging across participant responses) within the relevant groups (e.g., unfamiliar vs familiar, or specific superordinate classes), unless stated otherwise.

#### 4.4.3 Bayesian classification

Our Bayesian classifier was used to calculate the posterior probability of each possible superordinate class given a set of labelled parts and select that with the highest posterior probability. The parts in question were those provided as labels by a participant on a given trial. As such, the classifier operated on a trial-wise basis, which had the benefit of allowing direct comparison with participant responses. To construct this classifier, we implemented the standard Bayesian inference formula:

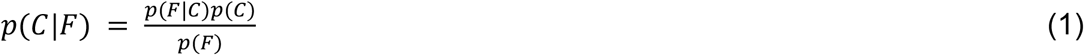

Where *C* represents a given class and *F* represents a given set of part labels. In the formula, *p(C*|*F)* represents the posterior, *p(F*|*C)* the likelihood, *p(C)* the prior, and *p(F)* the marginal.

To calculate *p(F*|*C)*, we need to incorporate the evidence provided by each part, *f*. For this, we combine their individual likelihoods, *p(f*|*C)*, by calculating their product:

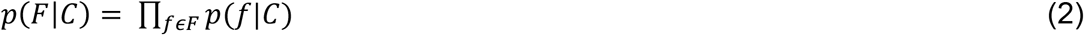

We can substitute this into Bayes’ rule, dropping the marginal as it is the same across all classes:

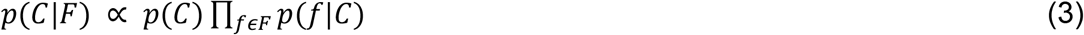

When calculating the combined part likelihood, *p(F*|*C)*, computing the product of probabilities results in very small numbers. To avoid issues related to floating-point underflow, we complete equation 3 in log space, i.e.:

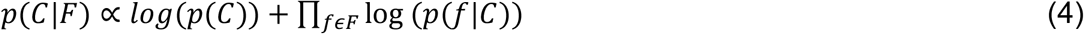

To avoid zero-values in our log calculations, we apply Laplace smoothing with an alpha value of 1 to our individual likelihood calculations. Once the posterior probabilities for all possible classes are calculated for a given trial and its relevant set of part labels, we subtract the maximum value. In doing so, we again avoid issues related to floating-point underflow resulting from subsequent exponentiation of large negative log values, while preserving the final probabilities. After applying the subtraction, we apply the exponent to all values and then normalise the values to provide the final probabilities (i.e., the posteriors) for each class.

For the classification, we simply take the class with the highest posterior probability value, *p(C*|*F)*. This process is replicated for each individual trial and its respective set of part labels. Therefore, we obtain a set of classification responses to the same images as participants and can compare our classifier’s performance to participants’ on a trial-wise basis.

#### 4.4.4 DreamSim embedding space and classification

To retrieve a similarity space for our drawings we used DreamSim which, compared to large vision models like DINO and CLIP, is human-aligned and bridges the gap between lower-level image features and broader categorical comparisons and has performance in sketch-based photo retrieval similar to openCLIP^28^. We obtained individual perceptual similarity vectors *v* by embedding each drawing into the DreamSim model and then normalised each vector:

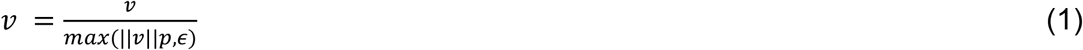

We then computed a similarity matrix of all drawings by calculating the cosine distance between the normalised embedding vectors of any two drawings:

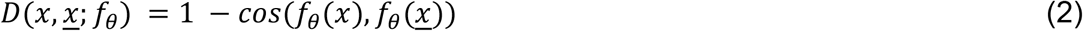

Where *f*_*θ*_ represents the embedding vector. We then transformed the similarity matrix by t-SNE to visualise similarity as distances between exemplars in a two-dimensional space. To compare the clustering of drawings in DreamSim embedding space for unfamiliar and familiar drawings, we obtained two class spread metrics by calculating within each class the mean pairwise distances between all exemplars and the mean distance to the cluster centre. Additionally, we calculated the pairwise distances of the centroids for each class, separately for familiar and unfamiliar drawings. This yielded two 10×10 distance matrices. To compare their values, we correlated the upper triangle (without the diagonal) values for both matrices.

