## Supplemental Materials for "How we draw and recognize things that don’t exist"

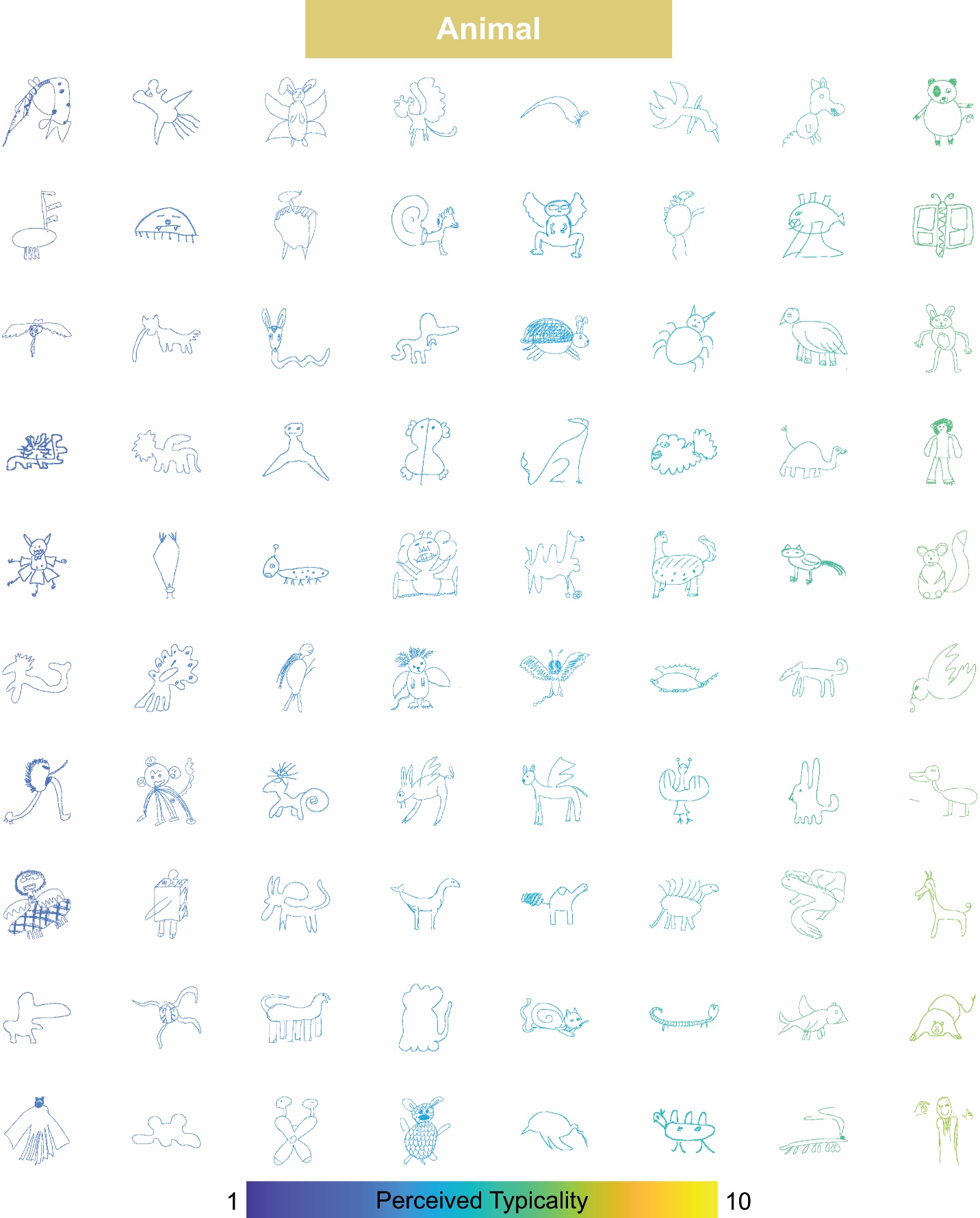

**Figure S1. Perceived typicality for unfamiliar animal drawings.** Drawings are arranged and colour-coded according to their average perceived typicality.

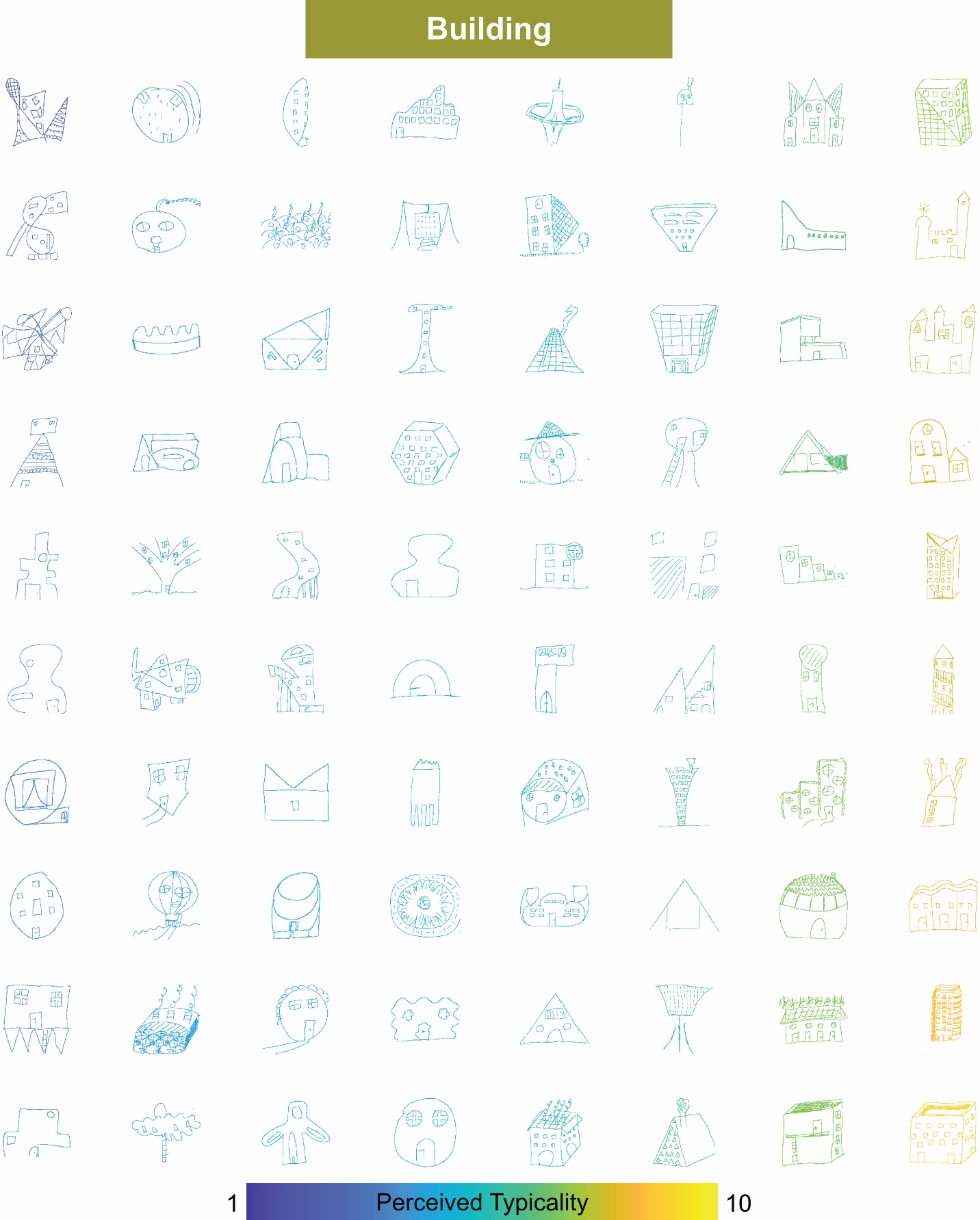

**Figure S2. Perceived typicality for unfamiliar building drawings.** Drawings are arranged and colour-coded according to their average perceived typicality.

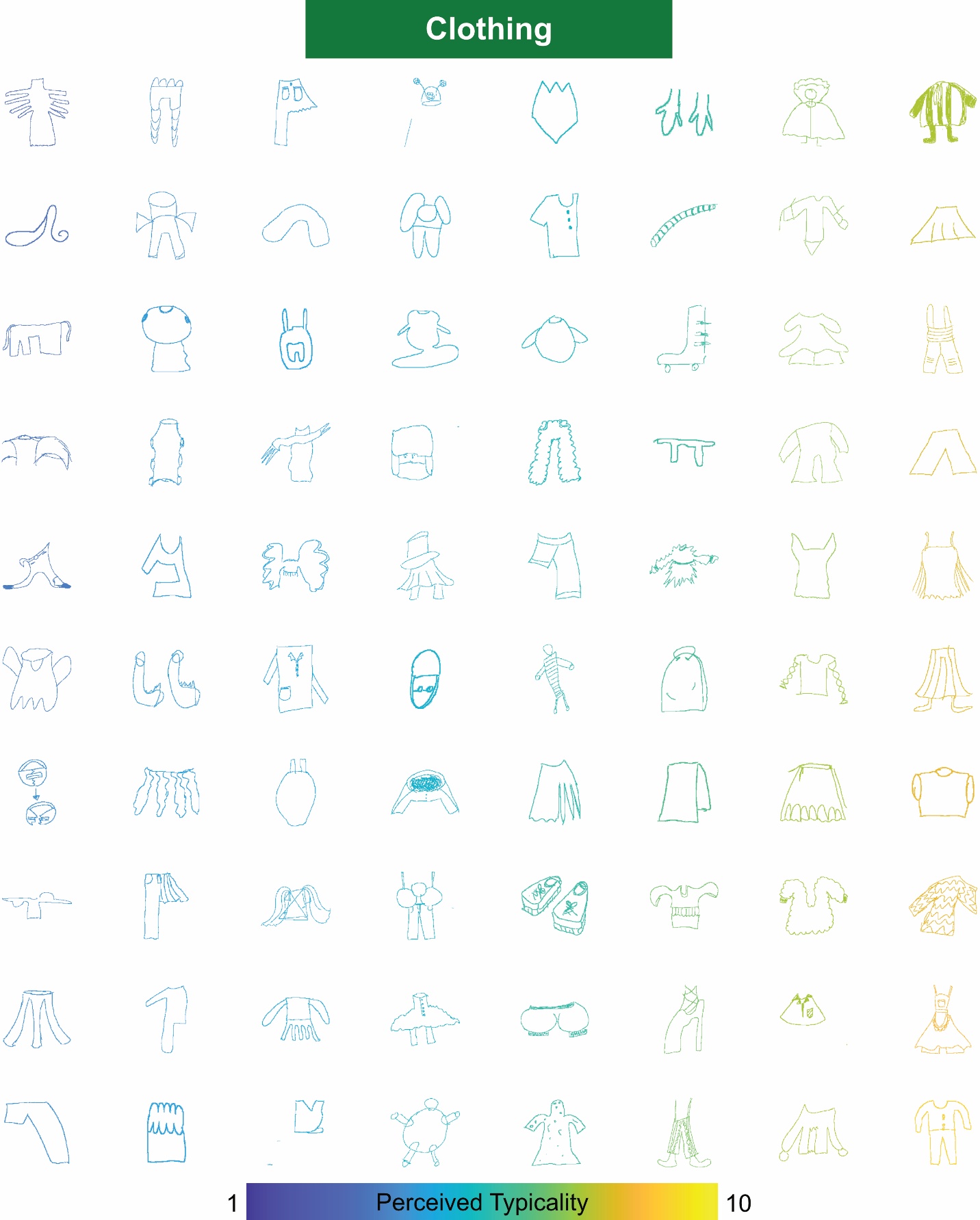

**Figure S3. Perceived typicality for unfamiliar clothing drawings.** Drawings are arranged and colour-coded according to their average perceived typicality.

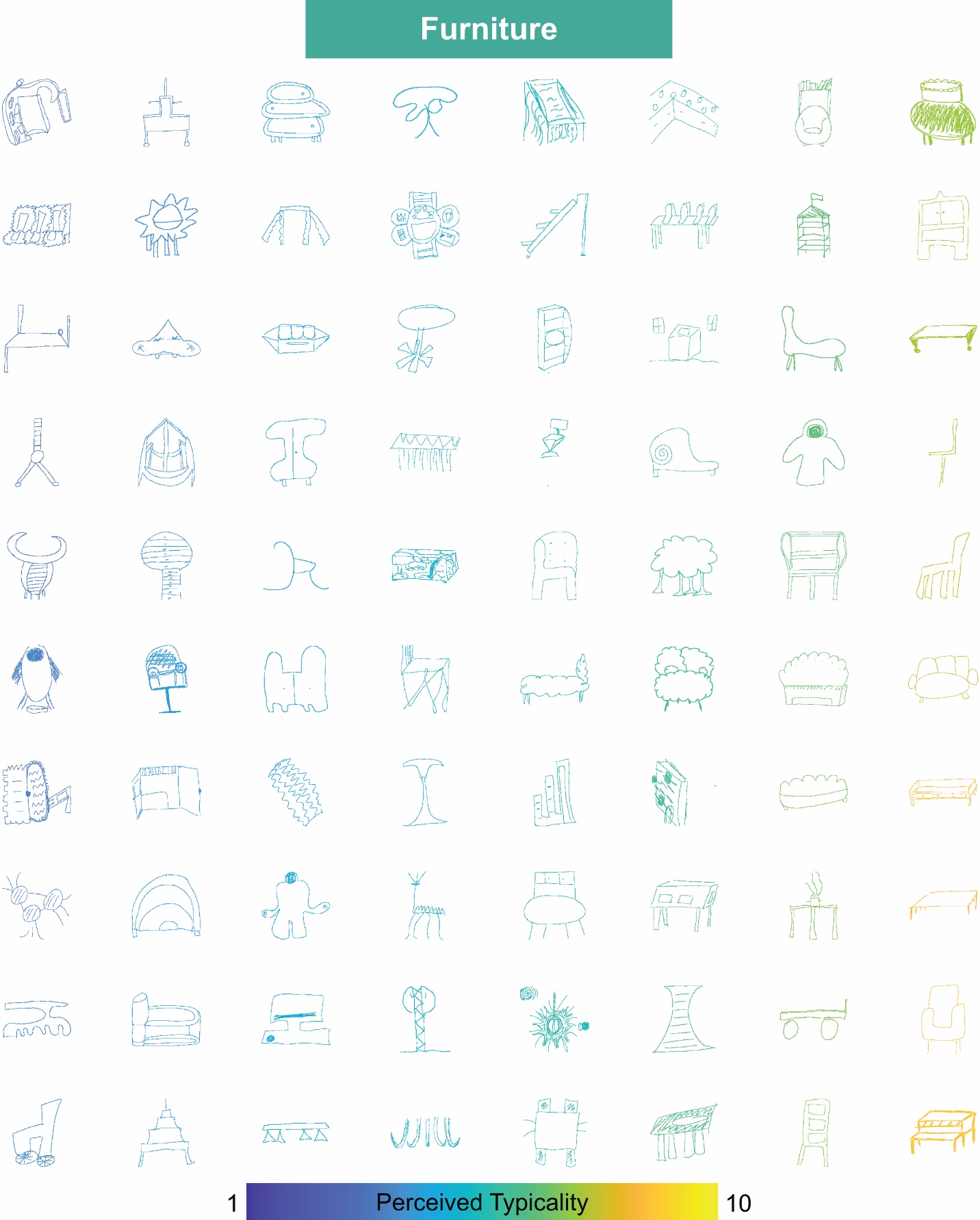

**Figure S4. Perceived typicality for unfamiliar furniture drawings.** Drawings are arranged and colour-coded according to their average perceived typicality.

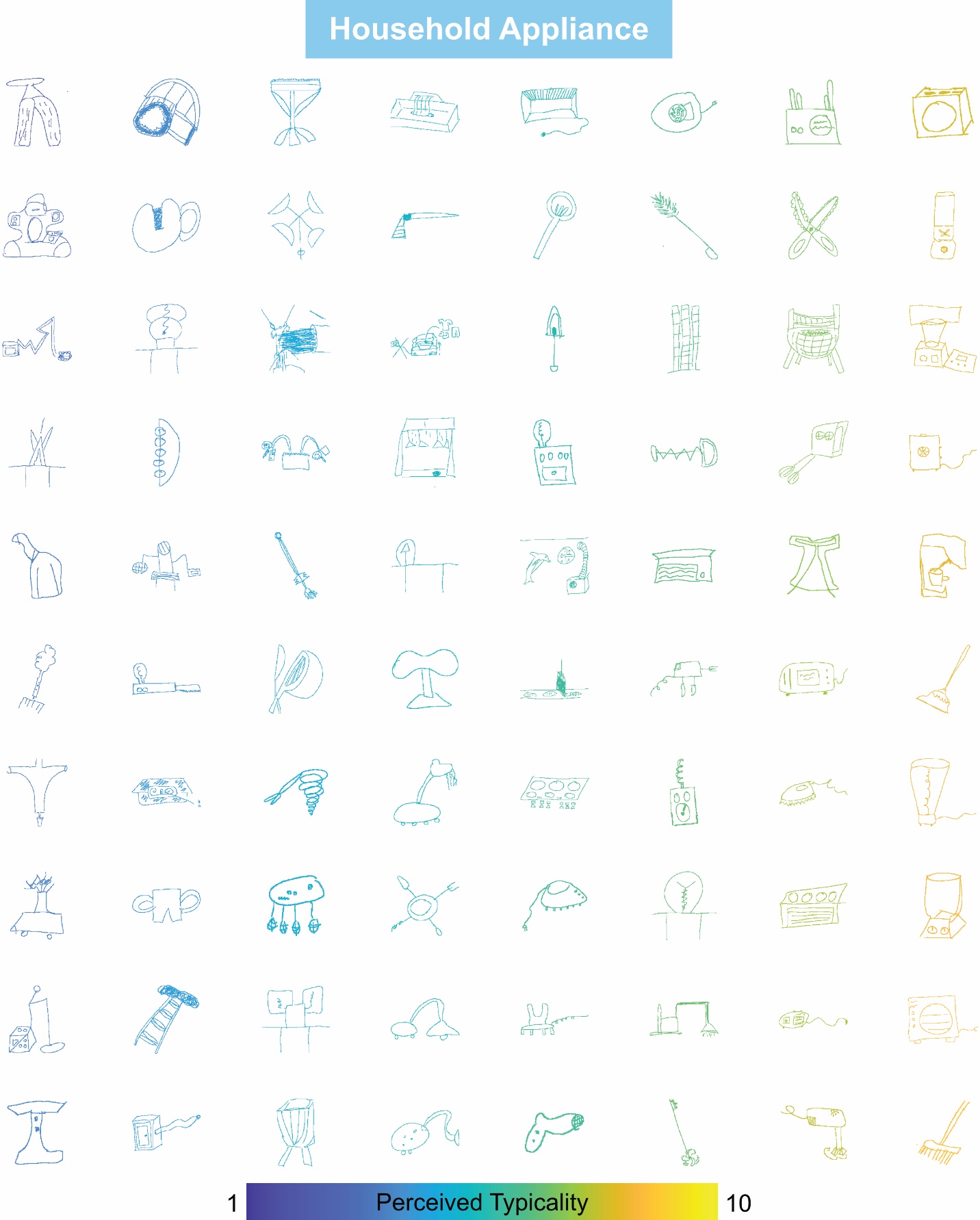

**Figure S5. Perceived typicality for unfamiliar appliance drawings.** Drawings are arranged and colour-coded according to their average perceived typicality.

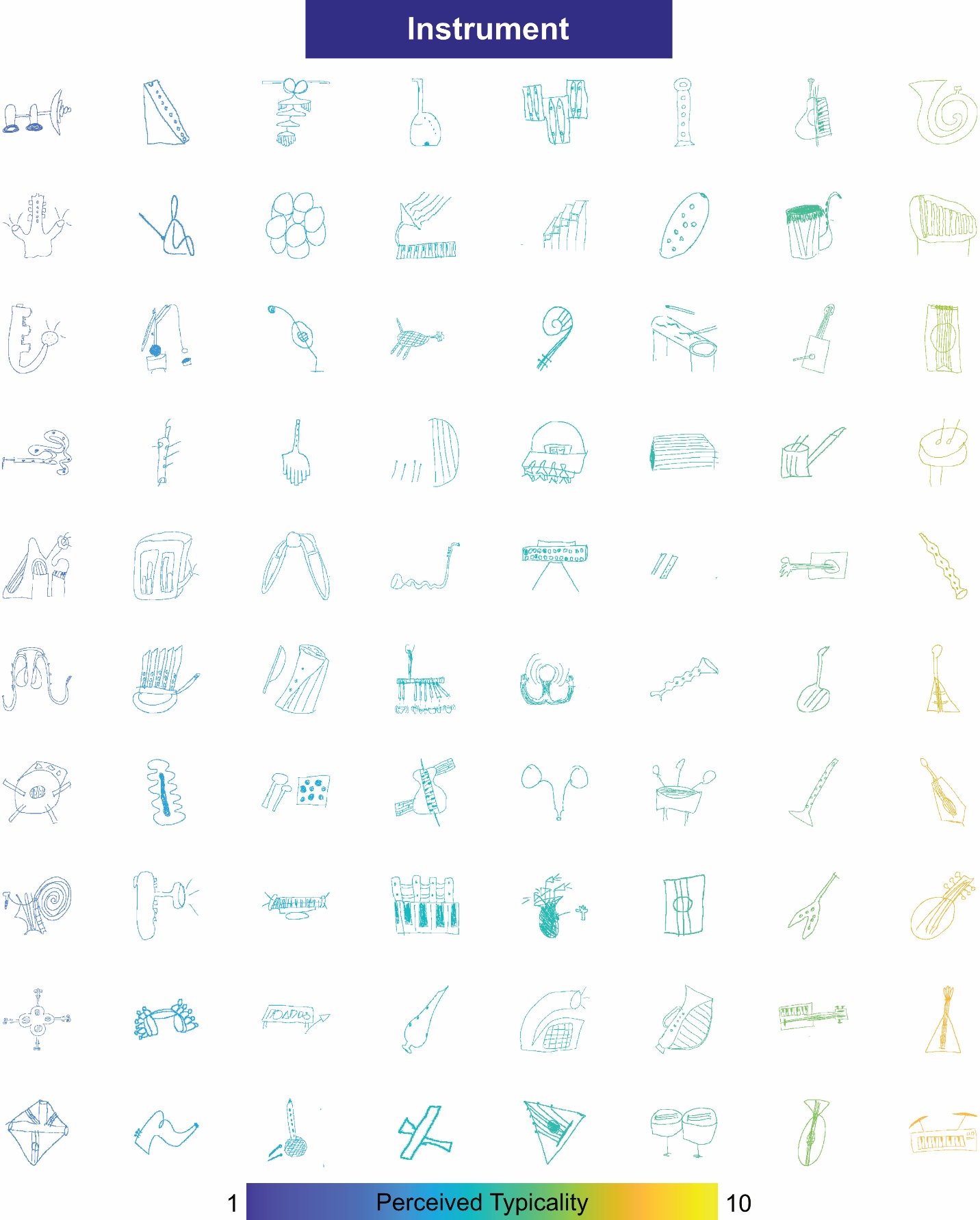

**Figure S6. Perceived typicality for unfamiliar instrument drawings.** Drawings are arranged and colour-coded according to their average perceived typicality.

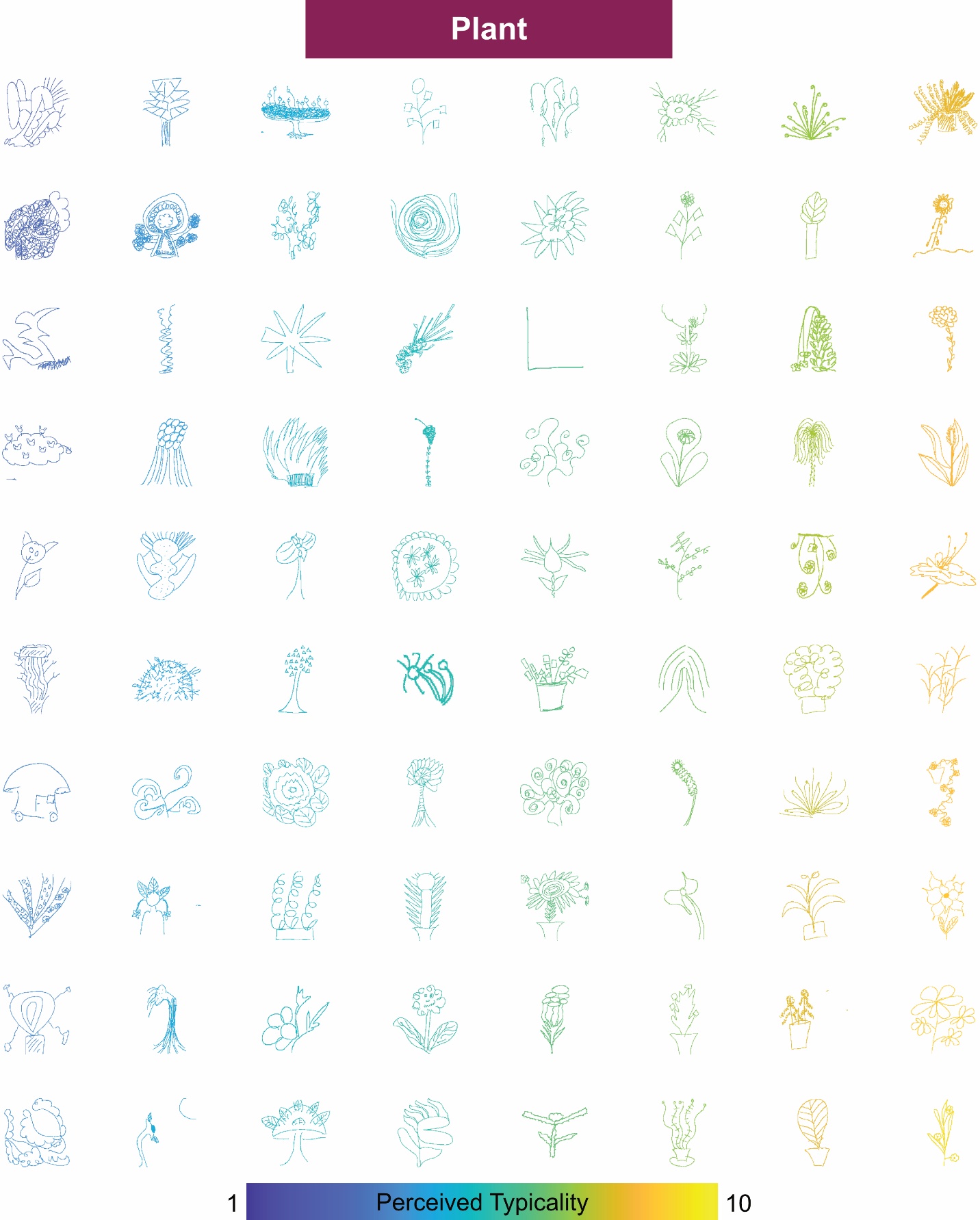

**Figure S7. Perceived typicality for unfamiliar plant drawings.** Drawings are arranged and colour-coded according to their average perceived typicality.

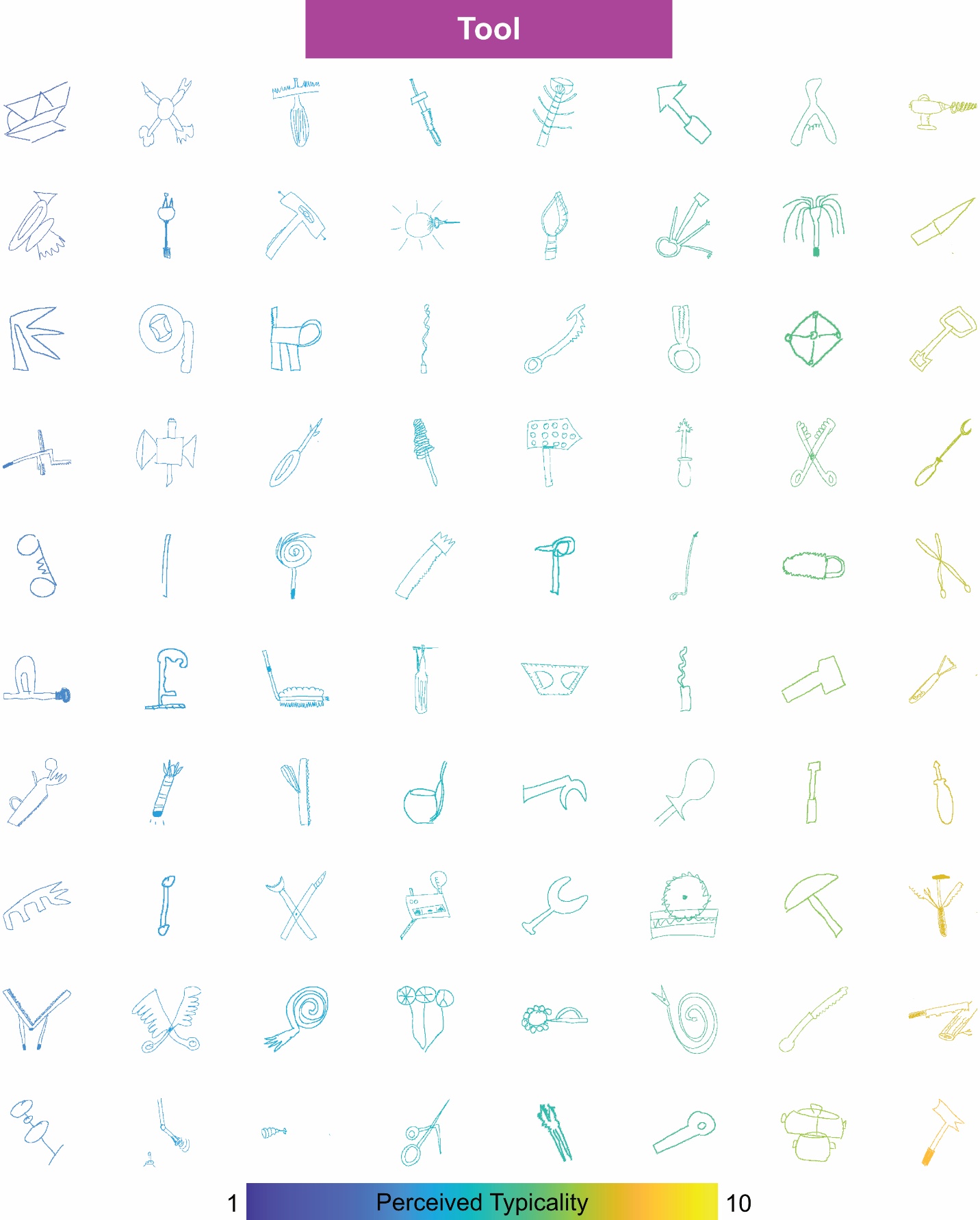

**Figure S8. Perceived typicality for unfamiliar tool drawings.** Drawings are arranged and colour-coded according to their average perceived typicality.

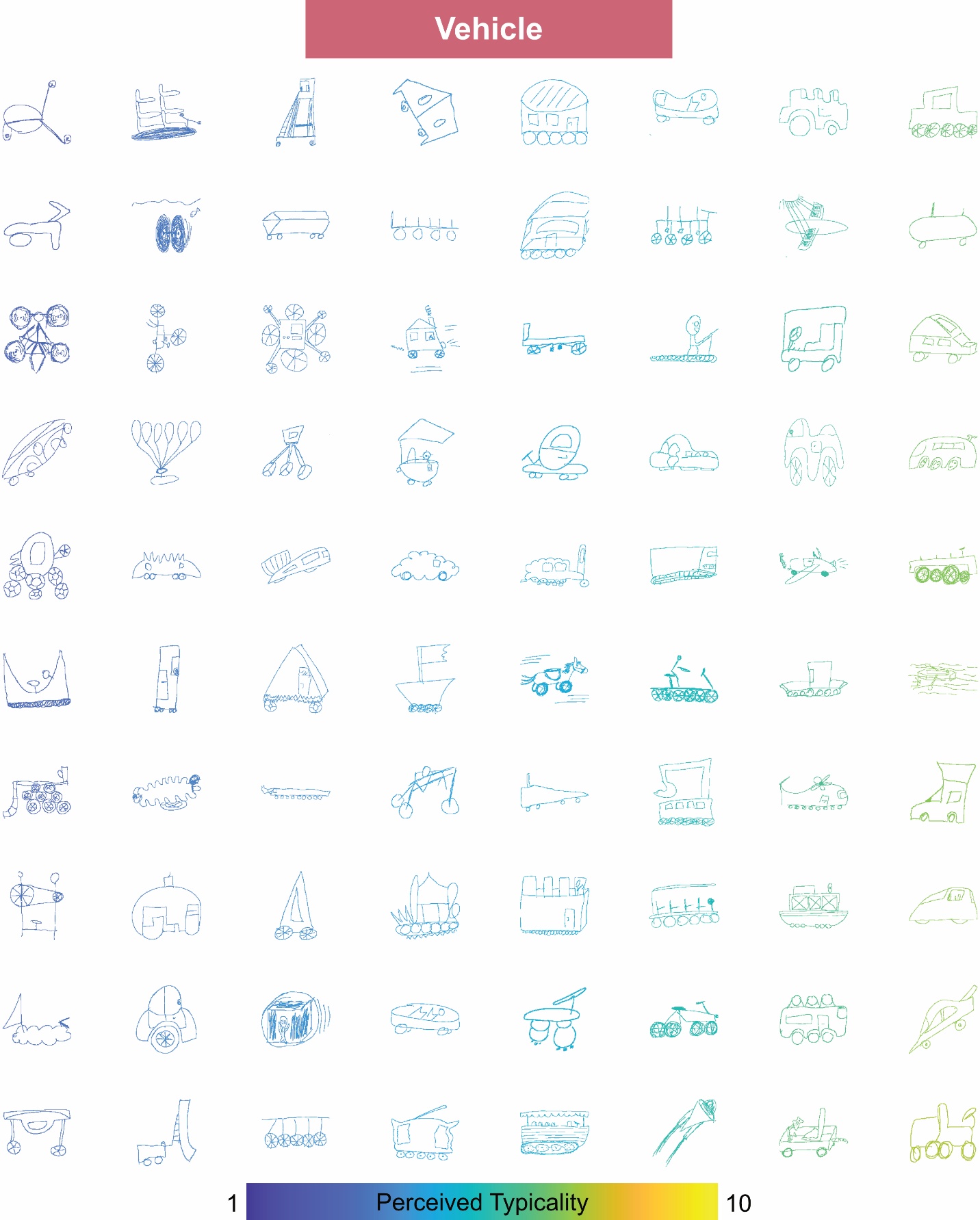

**Figure S9. Perceived typicality for unfamiliar vehicle drawings.** Drawings are arranged and colour-coded according to their average perceived typicality.

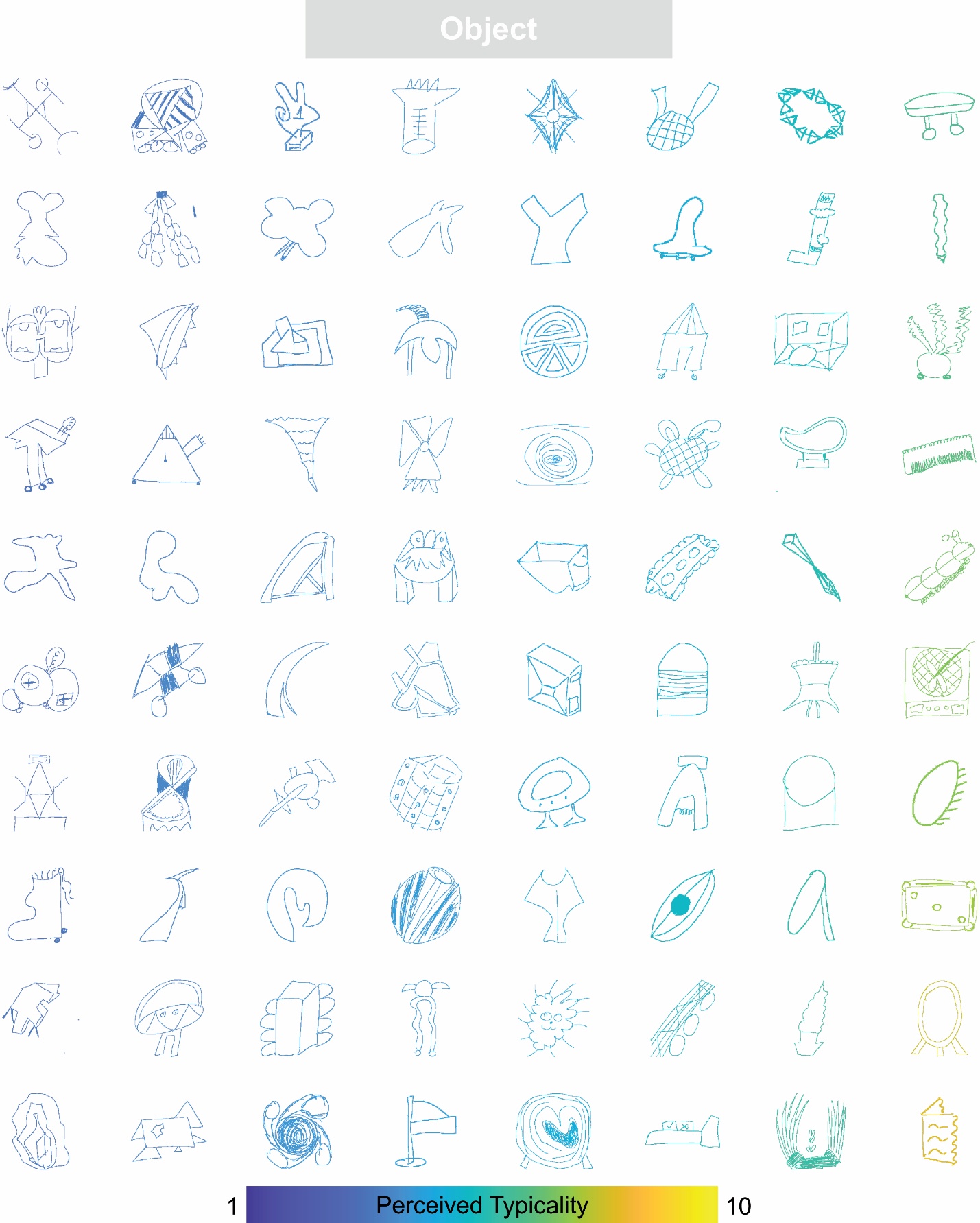

**Figure S10. Perceived typicality for unfamiliar object drawings.** Drawings are arranged and colour-coded according to their average perceived typicality.

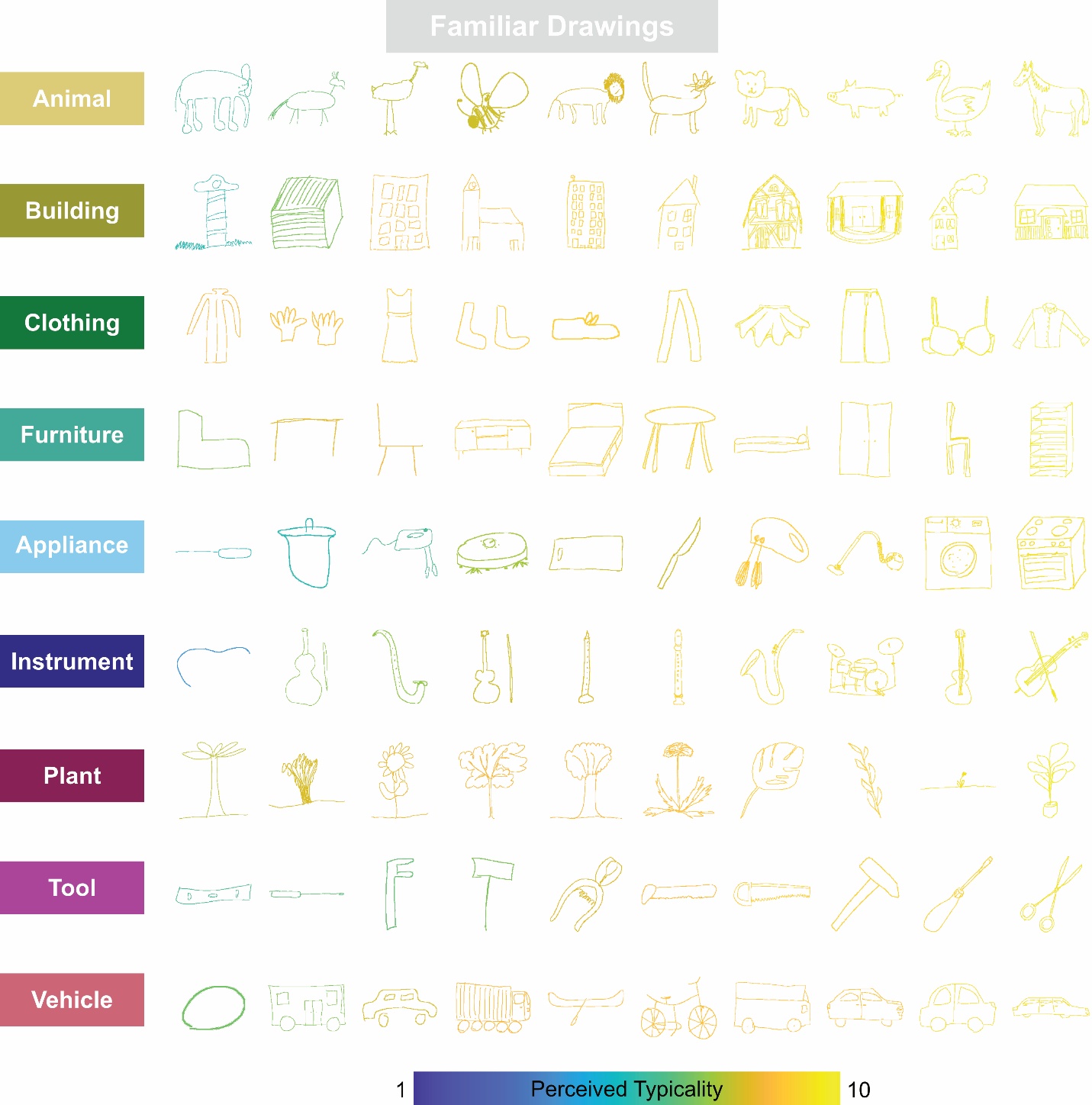

**Figure S11. Perceived typicality for all familiar drawings.** Drawings are arranged in rows according to their superordinate class, and colour-coded according to their average perceived typicality.

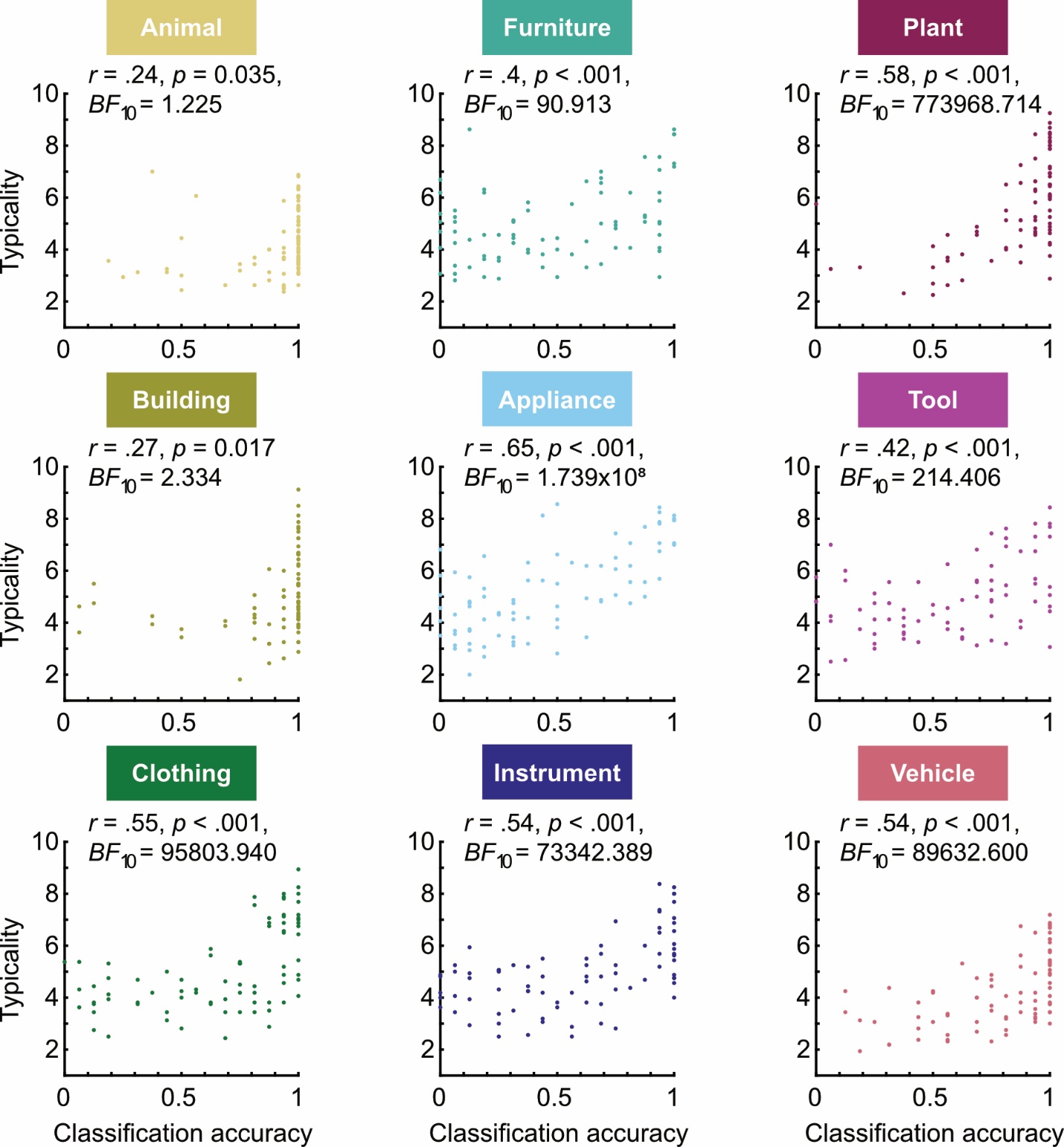

**Figure S12. Individual class correlations between typicality ratings and classification accuracy for unfamiliar drawings.** Each plot represents one superordinate class (according to the class cued to generate the drawings), with each datapoint representing an individual drawing. Included are relevant *r*, *p*, and *BF* values.

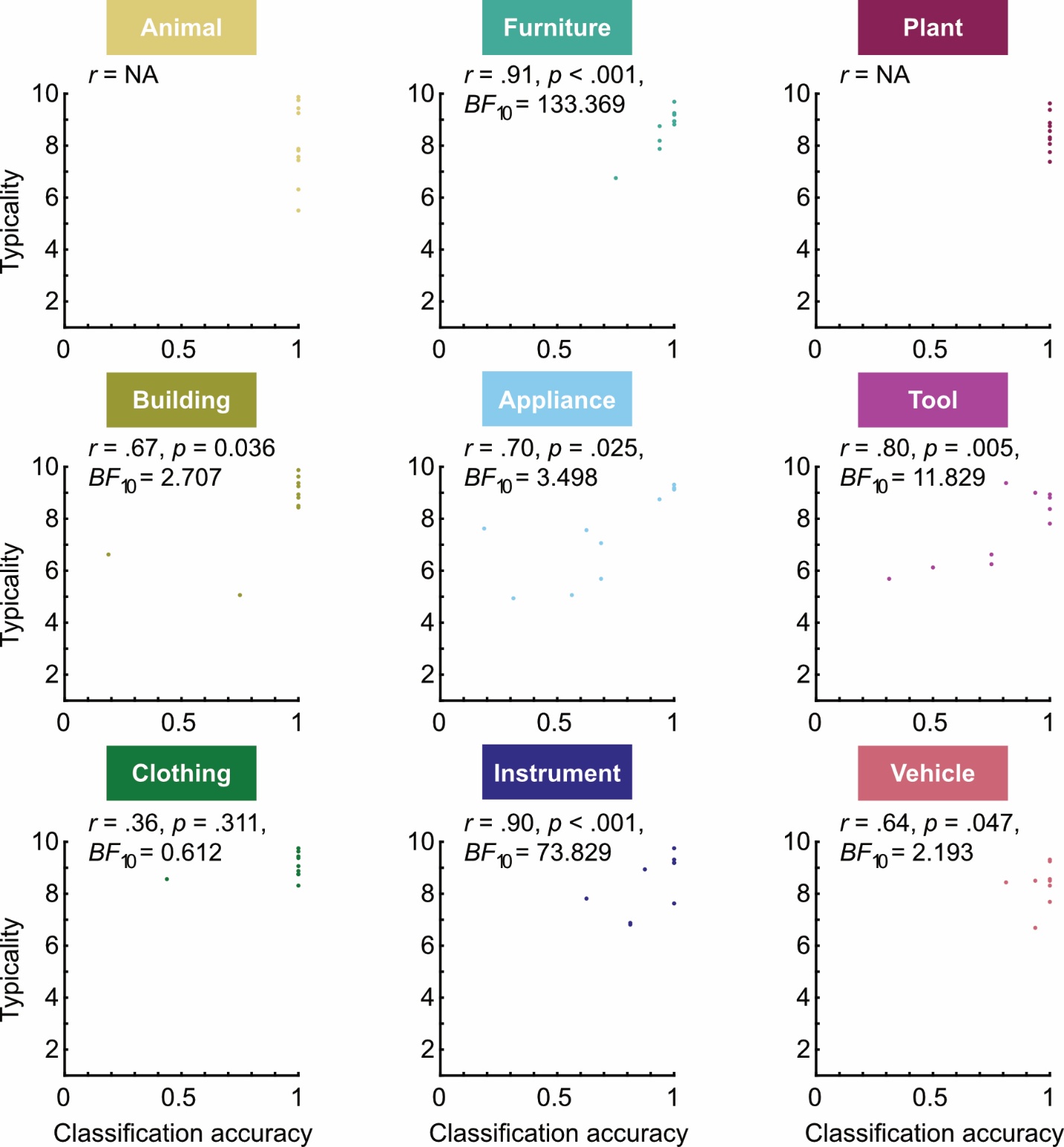

**Figure S13. Individual class correlations between typicality ratings and classification accuracy for familiar drawings.** Each plot represents one superordinate class (according to the class cued to generate the drawings), with each datapoint representing an individual drawing. Included are relevant *r*, *p*, and *BF* values. Note, we encourage caution when interpreting these relationships for familiar drawings due to the lower image sample size, and general skewing towards high classification accuracy.

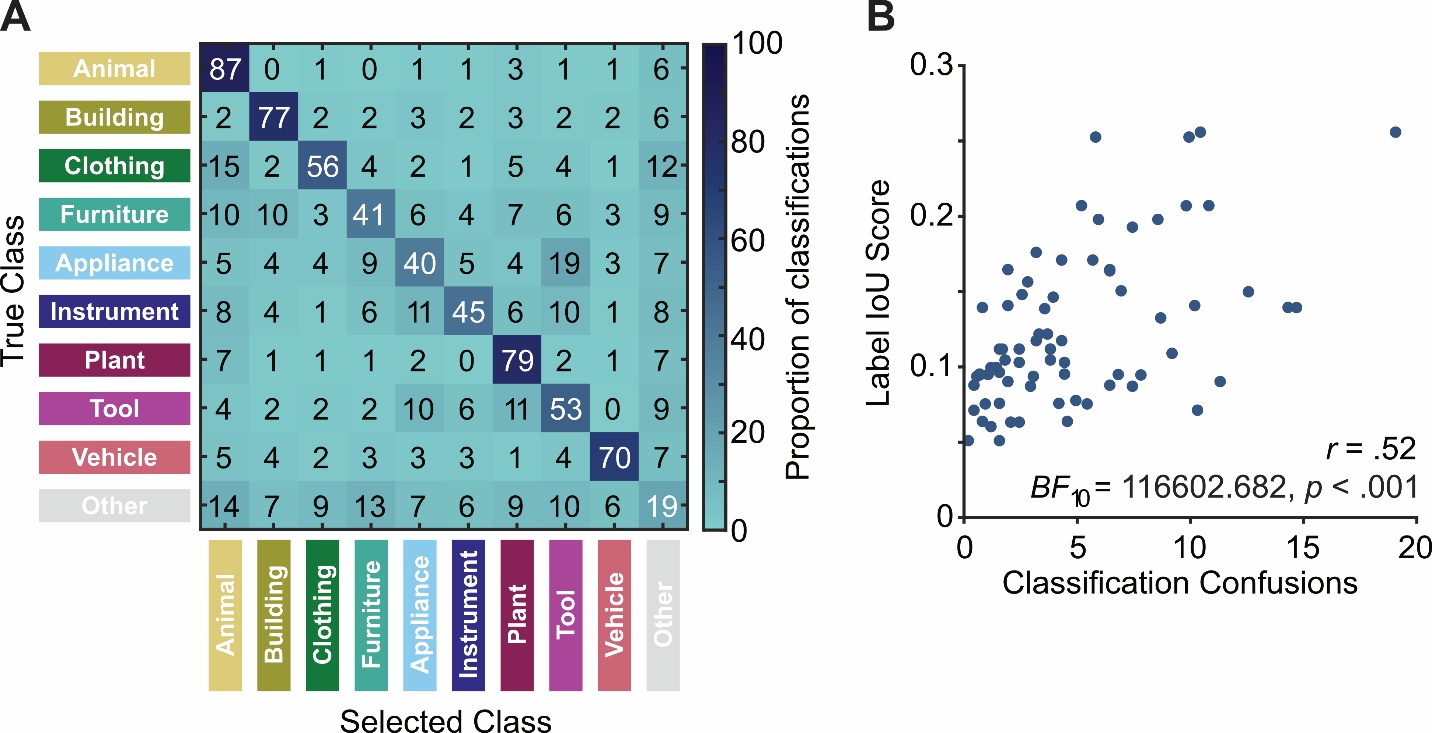

**Figure S14. Relating IoU with classification confusions for unfamiliar drawings.** To investigate the relationship between label distinctiveness and classification confusions, we look at the IoU score for each superordinate class pairing (per Fig. 6), and correlate this with the number of classification confusions specific to that pairing. **A)** Matrix representing the classification confusions made for images belonging to each superordinate class. Values represent the proportion of trials corresponding to the relevant superordinate class. For example, if we take the top row of panel A, we can calculate the number of times participants mis-classified the drawings of animals for a given alternative superordinate class. **B)** Here, we plot the classification confusion against the IoU value for the labels corresponding to the same position in the matrix (per Fig. 6). We find very strong evidence for a moderate positive correlation (r = .52, Bayesian correlation BF_10_ = 113503.515, p < .001), demonstrating more classification confusions when the IoU score is greater.

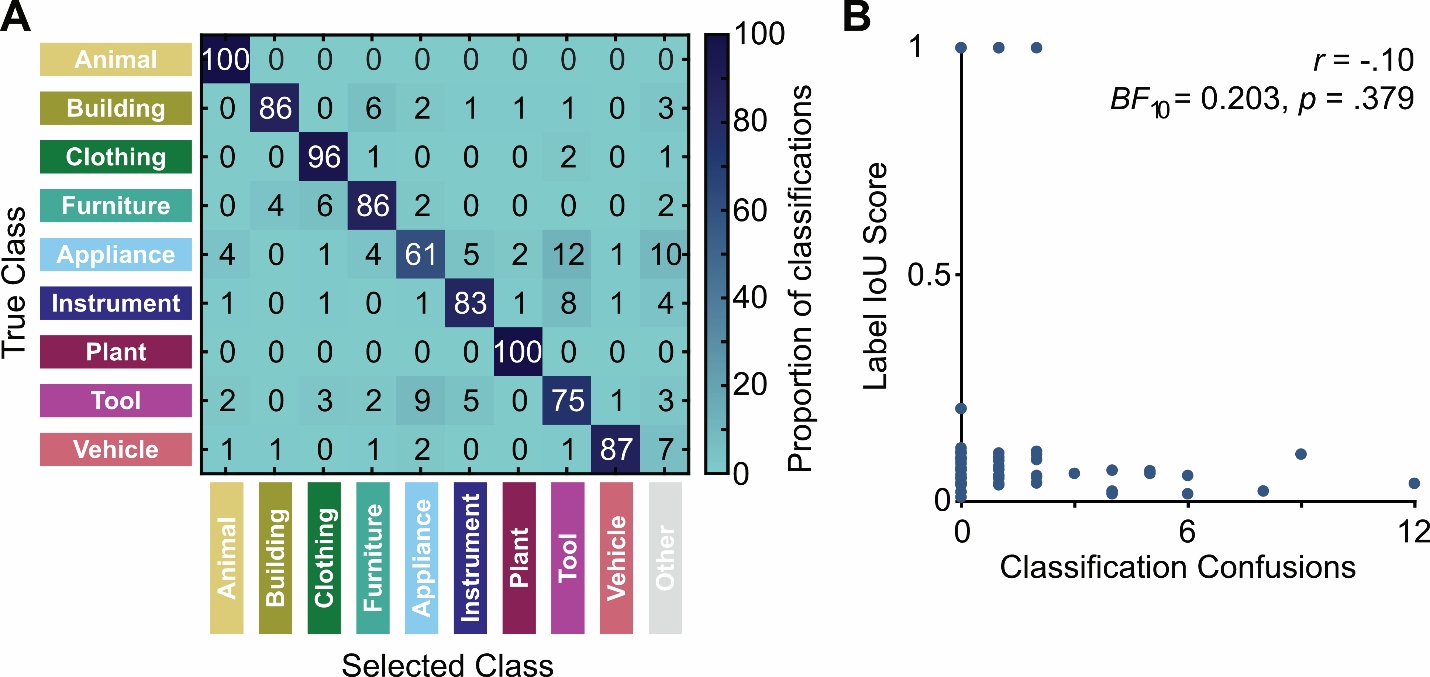

**Figure S15. Relating IoU with classification confusions for familiar drawings.** To investigate the relationship between label distinctiveness and classification confusions, we look at the IoU score for each superordinate class pairing (per Fig. 6), and correlate this with the number of classification confusions specific to that pairing. **A)** Matrix representing the classification confusions made for images belonging to each superordinate class. Values represent the proportion of trials corresponding to the relevant superordinate class. For example, if we take the top row of panel A, we can calculate the number of times participants mis-classified the drawings of animals for a given alternative superordinate class. Note, that object is not present as a True Class, as there were no ‘true’ familiar Object drawings tested. Nonetheless, familiar drawings could still be classified as other by participants, hence it being present as a potential Selected Class. **B)** Here, we plot the classification confusion against the IoU value for the labels corresponding to the same position in the matrix (per Fig. 6). We find moderate evidence in favour of there being no correlation (r = -.10, Bayesian correlation BF_10_ = 0.203, p = .379), likely due to heavy skewing of classifications resulting in very few classification confusions.

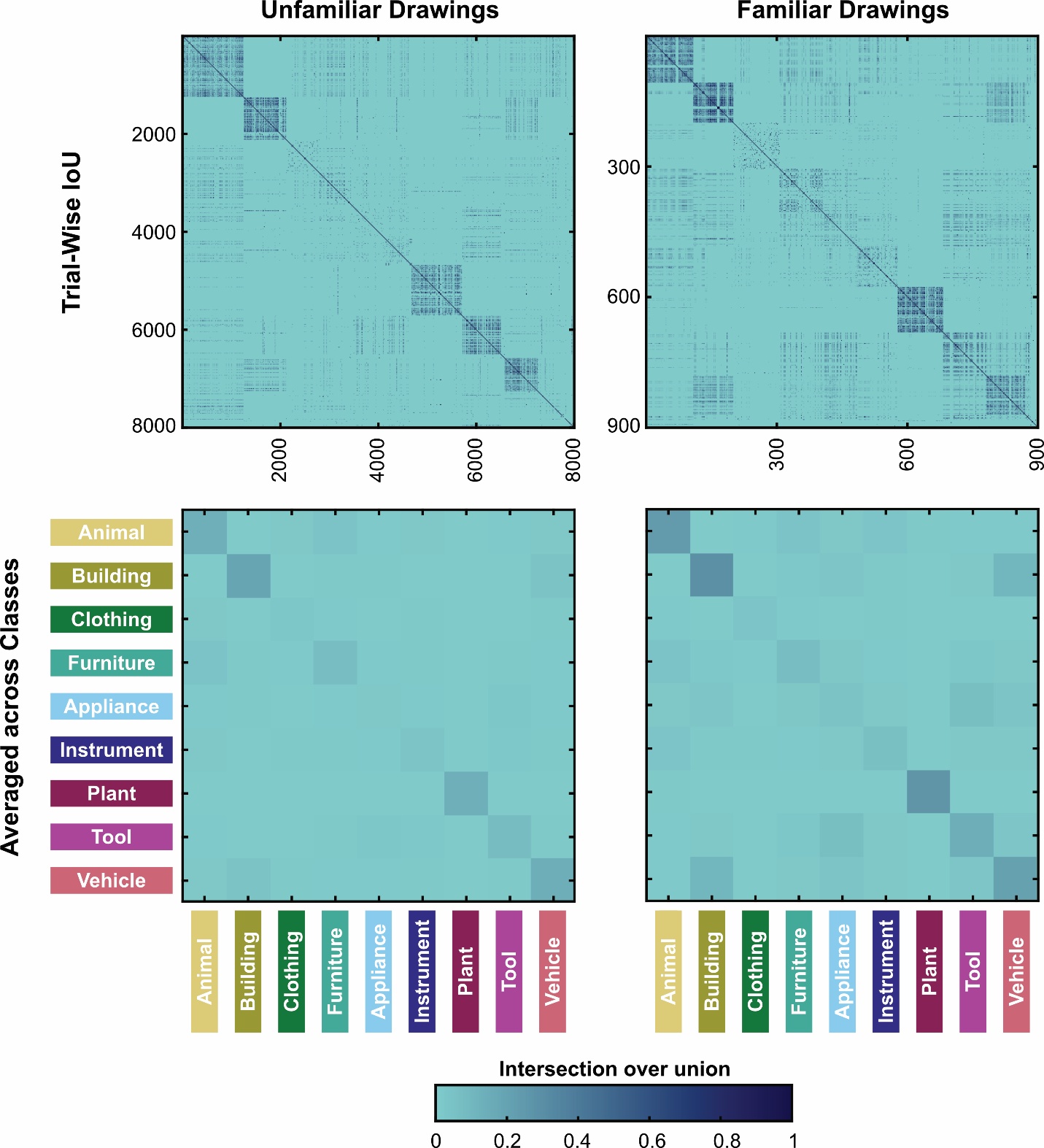

**Figure S16. Trial-wise intersection over union analysis.** We calculated the intersection over union value for labels provided to drawings on a trial-wise level (top row), completing all possible comparisons for all 8000 unfamiliar drawing trials (left) and all 900 familiar drawing trials (right). Trials are ordered based on the class assigned by participants in Experiment 3 before they provided the labels. We then calculate the average IoU value across trials within each class, and re-plot these average values (bottom row).

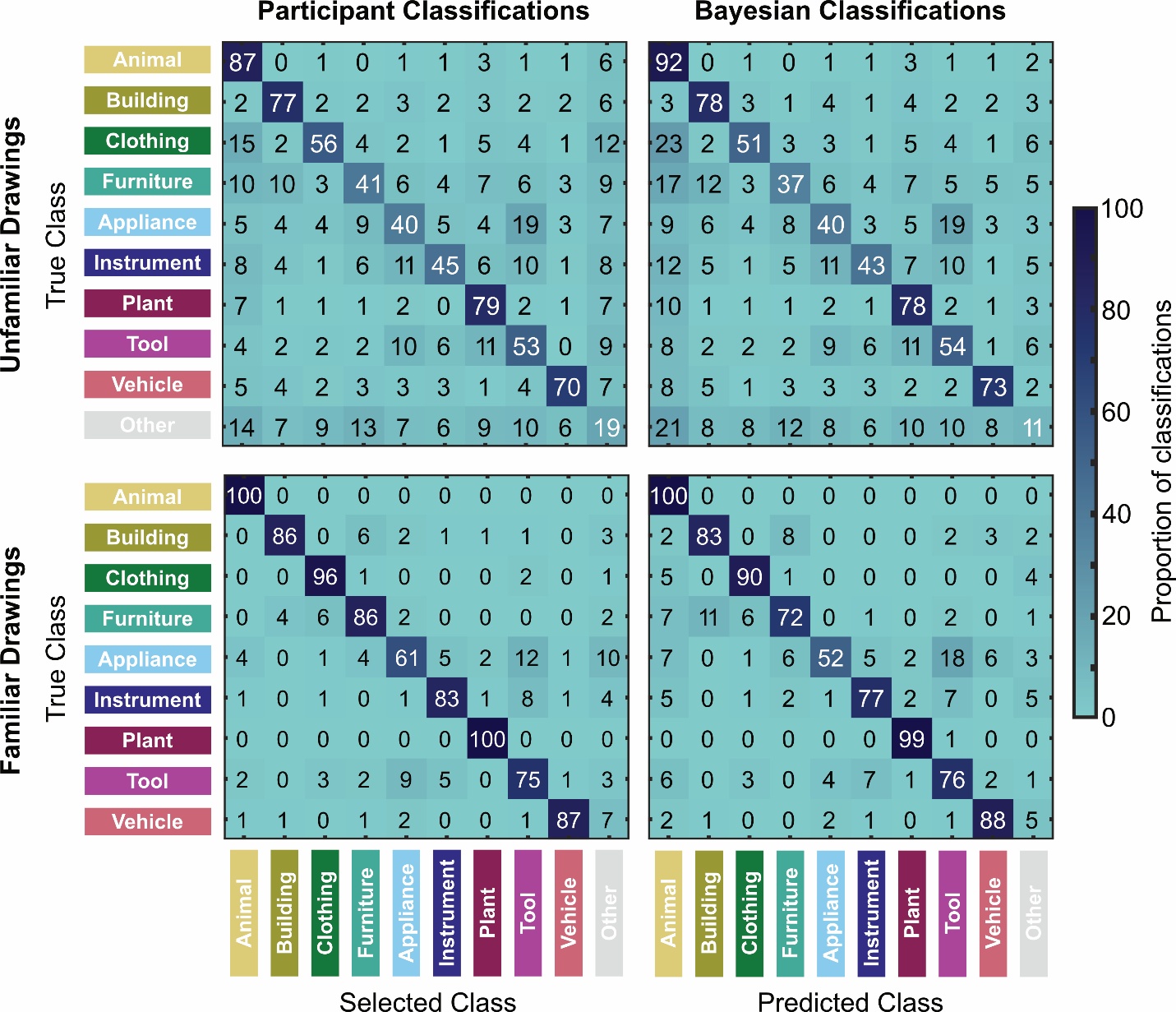

**Figure S17. Observer vs Bayesian classifier confusions for unfamiliar and familiar drawings.** Matrices representing the classification confusions made by human observers (left column) and a Bayesian classifier (right column) for images belonging to each superordinate class, for both unfamiliar (top row) and familiar (bottom row). Values represent the proportion of trials corresponding to the relevant superordinate class/familiarity condition. Note, for familiar drawings Other is not present as a True Class, as there were no ‘true’ other familiar drawings tested. Nonetheless, familiar drawings could still be classified as other by participants and the Bayesian classifier, hence it being present as a potential Selected Class.

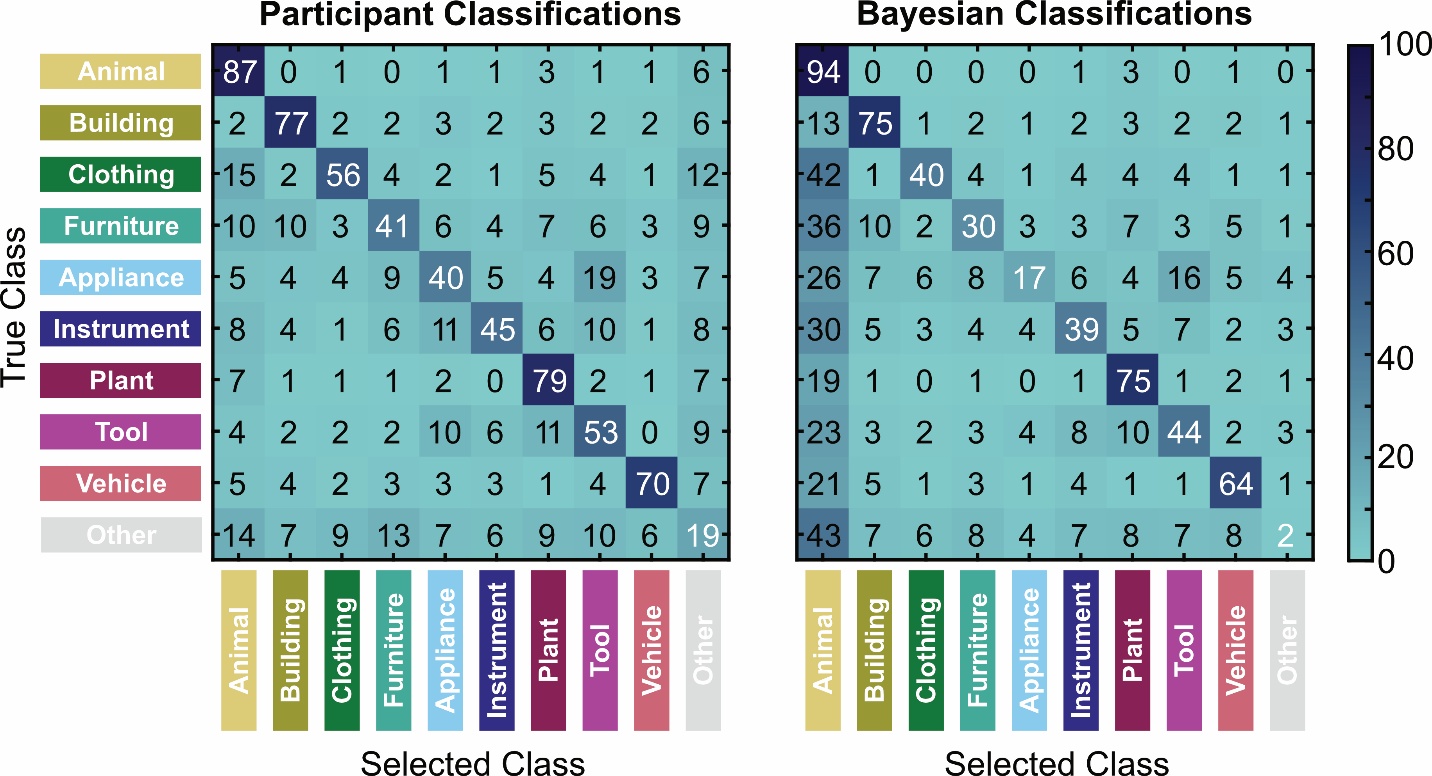

**Figure S18. Observer vs Bayesian classifier confusions for unfamiliar drawings, based on familiar priors.** Matrices representing the unfamiliar drawing classification confusions made by human observers (left) and a Bayesian classifier (right) which bases its priors on familiar drawing information. Values represent the proportion of trials corresponding to the relevant superordinate class/familiarity condition. As not all part labels present for unfamiliar drawings were also present for familiar drawings, some unfamiliar drawings had no likelihood information to derive from their parts. In this case, the classification is driven by the class priors, which are based on the number of drawings classified into the various superordinate classes by participants. Such ratings resulted in a higher count of animal objects than any other class, meaning that in the case where no likelihood information can be derived from the parts, the classifier selected animal. As a result, there is a bias for animal classifications, which can be seen by the higher classification confusions attributed to animals (left-most column for Bayesian classification matrix).
